# Glia-derived secretory fatty acid binding protein Obp44a regulates lipid storage and efflux in the developing *Drosophila* brain

**DOI:** 10.1101/2024.04.10.588417

**Authors:** Jun Yin, Hsueh-Ling Chen, Anna Grigsby-Brown, Yi He, Myriam L. Cotten, Jacob Short, Aidan Dermady, Jingce Lei, Mary Gibbs, Ethan S. Cheng, Dean Zhang, Caixia Long, Lele Xu, Tiffany Zhong, Rinat Abzalimov, Mariam Haider, Rong Sun, Ye He, Qiangjun Zhou, Nico Tjandra, Quan Yuan

## Abstract

Glia derived secretory factors play diverse roles in supporting the development, physiology, and stress responses of the central nervous system (CNS). Through transcriptomics and imaging analyses, we have identified Obp44a as one of the most abundantly produced secretory proteins from *Drosophila* CNS glia. Protein structure homology modeling and Nuclear Magnetic Resonance (NMR) experiments reveal Obp44a as a fatty acid binding protein (FABP) with a high affinity towards long-chain fatty acids in both native and oxidized forms. Further analyses demonstrate that Obp44a effectively infiltrates the neuropil, traffics between neuron and glia, and is secreted into hemolymph, acting as a lipid chaperone and scavenger to regulate lipid and redox homeostasis in the developing brain. In agreement with this essential role, deficiency of Obp44a leads to anatomical and behavioral deficits in adult animals and elevated oxidized lipid levels. Collectively, our findings unveil the crucial involvement of a noncanonical lipid chaperone to shuttle fatty acids within and outside the brain, as needed to maintain a healthy brain lipid environment. These findings could inspire the design of novel approaches to restore lipid homeostasis that is dysregulated in CNS diseases.

## Introduction

Lipids are essential components of the brain, serving as vital substrates for membrane formation, signal transduction, as well as energy supply and storage. Not only is the brain a lipid rich organ but its lipid composition is remarkably complex. Many brain lipid species are exclusively found in neural tissues, suggesting specialized lipid requirements for brain functions (Bozek et al., 2015; Fitzner et al., 2020; Vaughen et al., 2022). How does the brain maintain the integrity and stability of its unique lipid environment? Animals have evolved several tightly regulated processes to address this challenge: the uptake of lipids from circulation into the brain via the blood-brain barrier (BBB), intra and intercellular lipid transport and storage, local lipid synthesis and metabolism, and lastly, the recycling and efflux of lipids out of the brain (Kadry et al., 2020; Rhea & Banks, 2021). In the central nervous system (CNS), these complex processes are often orchestrated through neuron-glia interactions (Barber & Raben, 2019; Chung et al., 2024; Liu et al., 2017). Dysregulation of these mechanisms is associated with neurological disorders and neurodegenerative diseases (Brandebura et al., 2023; Caldwell et al., 2022; Goodman & Bellen, 2022), highlighting the importance of maintaining a stable lipid environment for optimal brain function (Valles & Barrantes, 2022).

Despite the importance of brain lipid homeostasis, the mechanisms underlying its regulation are challenging to investigate. While general brain lipid homeostatic strategies are utilized across the entire animal kingdom, the CNS lipid environment of individual species is strongly influenced by factors such as diet, environment, lifestyle, and developmental progression (Carvalho et al., 2012). The heterogeneous distribution of lipids is another significant hindrance to brain lipid research. Recent lipodomics and metabolomics analyses of the mammalian brain have revealed lipid profiles that are regulated in a regional and cell-specific way throughout development (Fitzner et al., 2020). Even though brain lipid heterogeneity is well documented, the regulatory mechanisms governing brain lipid trafficking and homeostasis remain largely unexplored.

In recent years, interest has grown to investigate a diverse array of molecular carriers of lipids, such as lipid chaperones and transporters (Chiapparino et al., 2016; Wong et al., 2019). These lipid transfer proteins share common structural features, often containing a hydrophobic pocket to bind, and thus mobilize hydrophobic lipids in the aqueous cytoplasm. Ultimately, they transfer lipids between cellular membranes and organelles, establishing temporal and spatially defined lipid distributions. Notably, these lipid transfer proteins can exhibit affinities that are specific to distinct lipid species, enabling them to adapt to various cellular contexts to perform their functional roles. Some well-known examples include phospholipid transfer proteins (PLTP) (Tall et al., 1985), sterol carrier protein 2 (SCP2) (Chanderbhan et al., 1982), Niemann-Pick C2 (NPC2) protein (McCauliff et al., 2011; Xu et al., 2008), and fatty acid binding proteins (FABPs) (Chiapparino et al., 2016; Ockner et al., 1972; Wong et al., 2019). Although lipid studies in the nervous system remain challenging, cellular and molecular understanding of lipid transfer proteins has been informative in terms of better understanding the basic biology and disease relevance of brain lipid regulation (Herz & Bock, 2002; Herz & Chen, 2006).

Among the molecules involved in brain lipid trafficking and metabolism, FABPs are lipid transfer proteins (LTPs) that act as specific molecular shuttles for fatty acids, playing a role in fatty acid oxidation, storage, and signaling pathways (Furuhashi et al., 2008). Broadly speaking, FABPs control the movement of fatty acids to various intracellular compartments and mediate the uptake of fatty acids from the extracellular environment into specific intracellular organelles such as the mitochondria, endoplasmic reticulum (ER) and peroxisome, where they are utilized for energy production and membrane synthesis. In the mammalian system, ten FABPs have been identified, each exhibiting unique patterns of tissue expression (Storch & Corsico, 2023). Of particular interest is the brain-specific FABP7, which is abundantly expressed in the embryonic mouse brain and later in the radial glia cells and astrocytes (Bennett et al., 1994; Ebrahimi et al., 2016). Intriguingly, FABP7 is upregulated in patients with Down syndrome and schizophrenia (Sanchez-Font et al., 2003; Watanabe et al., 2007). Recent studies also suggest that FABP7-deficient mice display altered emotional and behavioral responses and disrupted sleep wake cycles (Owada et al., 2006; Shimamoto et al., 2014). However, the impact of FABPs on the cell biology and lipid metabolism in CNS remains to be fully characterized.

While much of the research on FABPs has been conducted in mammals, *Drosophila* also has a FABP homolog that shares structural and functional similarities with its mammalian counterparts (Jang et al., 2022). *Drosophila* FABP (dFABP) has been shown to locate in the cortex and surface glia of the larval brain, where it regulates the formation of lipid droplets, a major source of lipid storage for the fly CNS (Kis et al., 2015). Unexpectedly, through transcriptomic analysis of astrocyte secretomes, we have identified a noncanonical FABP, Odorant binding protein 44a (Obp44a). As one of the most abundantly expressed secreted protein by astrocytes in both larval and adult brains, Obp44a exhibits affinity for long-chain fatty acids and exerts a broad impact on fatty acid trafficking and storage in the developing fly brain. Remarkably, Obp44a traffics within the brain and is secreted into the hemolymph, thereby facilitating the efflux of oxidized lipids out of the brain during development and oxidative stress conditions. Thus, OBP44a acts not only intracellularly, as known FABPs do, but also extracellularly. Collectively, the discovery of this new type of FABPs, along with the direct visualization of its co-trafficking with fatty acids, offers both functional characterization and mechanistic insights into the impact of FABP on brain lipid metabolism. It suggests the existence of a new class of FABPs as an innovative strategy adopted by insects to support brain development via effective transport of intact and oxidized fatty acids not only within but also outside the brain.

## Results

### Obp44a is a secreted molecule produced by the *Drosophila* CNS glia

Glia-derived secreted proteins serve key functions in neuron-glia interactions but have not been fully evaluated in the *Drosophila* system (Allen & Eroglu, 2017; Bittern et al., 2021; Farhy-Tselnicker & Allen, 2018; Liu et al., 2017). To analyze the astrocyte secretome, we performed in silico analysis using multiple published RNAseq datasets. They are generated either by single cell RNA sequencing (Sc-seq) or bulk RNA sequencing of FACS-sorted astrocytes at three different developmental stages (Brunet Avalos et al., 2019; Huang et al., 2015; Ravenscroft et al., 2020). Among the highly enriched transcripts in astrocytes, we specifically searched for secreted proteins identified through *Drosophila* extracellular domain database, FlyXCDB (Pei et al., 2018), which contains 1709 secreted proteins uncovered by robust computational predictions and manual curations.

The Sc-RNAseq dataset collected from the first instar larval brain contains 44 astrocytes, which are characterized by high enrichment of *alrm* transcripts (Brunet Avalos et al., 2019). Transcripts of 258 secreted proteins were detected in these astrocytes. Among the top highly enriched transcripts, Odorant Binding Protein 44a (Obp44a) is the most abundantly transcribed gene in first instar larval astrocytes (table S1, Fig. 1A, B). Subsequent investigations, using astrocyte-specific bulk RNAseq datasets collected from the third instar larval (L3) and adult stages (Huang et al., 2015), further reveal transcripts of 176 and 29 secreted proteins (table S2 and S3), respectively. Notably, Obp44a maintains high expression during brain development (fig. S1). Among the 13 secreted molecules that express across all developmental stages in astrocytes, Obp44a also exhibits the highest expression level (Fig.1C and fig. S1C, table S4).

**Figure 1.**
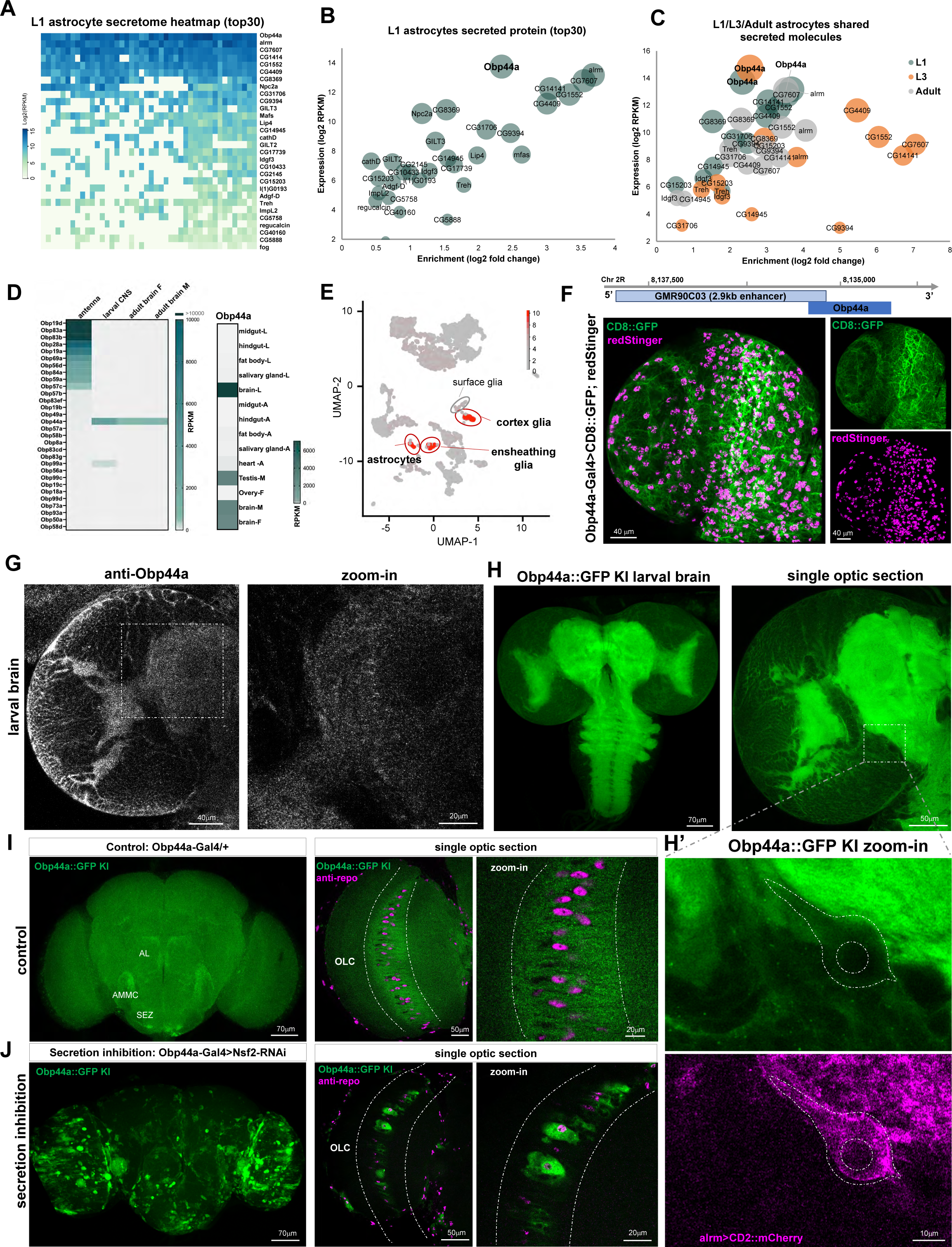
Obp44a is a one of the most abundant secretory proteins produced by *Drosophila* CNS glia. (A) Heatmap displaying the top 30 highly expressed and enriched secreted proteins in L1 astrocytes. Obp44a ranks as the most highly expressed secreted protein. (B) Expression and enrichment analysis of the top 30 secreted proteins in L1 astrocytes. (C) Expression and enrichment analysis of 13 common secreted molecules across three developmental stages (L1, L3, and adult), during which Obp44a consistently shows high expression levels. (D) Left: Expression heatmap presenting the top 30 expressed members of the Obp family in antenna, larval and adult brains. Obp44a shows high expression in the brain rather than the antenna, unlike the other classical odorant binding proteins. Right: Expression heatmap depicting Obp44a’s distribution across tissues, with prominent expression in the brain and testis. (E) Cell atlas of the L1 larval brain highlighting Obp44a’s high expression in three glial cell types: astrocytes, cortex glia, and ensheathing glia. (F) Obp44a enhancer labels a subpopulation of CNS glia. (Top) Obp44a-Gal4 is generated using a 2.9 Kb enhancer sequence located upstream of the transcriptional unit. (Bottom) The broad distribution of Obp44a in the larval brain is detected by Obp44a-Gal4 driven CD8::GFP (labels cell surface) or redStinger (labels nuclei). (G) Antibody staining revealing the endogenous localization Obp44a protein in the L3 larval brain, with a notable presence in the neuropil regions. (H) An Obp44a::GFP knock-in line is generated to directly visualize the distribution of Obp44a in the fly brain. Similar to anti-Obp44a staining, the signal prominently localizes in the neuropil region. (H’) Colocalization with astrocytes labeled by Alrm>CD2::mCherry shows a low level of Obp44a in astrocyte soma, consistent with the efficient secretion of the protein from astrocytes. (I) Obp44a::GFP shows a wide and diffused distribution in the adult brain. A single optic section of the optic lobe region demonstrates the Obp44a’s expression in the optic lobe chiasm (OLC) glia. (J) Blocking Obp44a secretion using Obp44a-Gal4 driven Nsf2 RNAi leads to concentrated GFP signal within glial soma, validating its glial production and secretion. Representative MIP or single optic sections of confocal images are shown. Scale bars are as indicated.

Additional transcriptome analysis indicates that, although Obp44a belongs to the odorant binding protein family, which facilitates the transport of hydrophobic odorant molecules, the expression of Obp44a in the olfactory sensory organ antennae is nearly undetectable (Fig. 1D) (Larter et al., 2016). Instead, Obp44a displays high expression levels in larval and adult brains, as well as in the male fly testis (Fig. 1D), shown by the ModEncode tissue specific RNAseq datasets (mod et al., 2010). To further investigate specific brain cell types responsible for Obp44a production, we analyzed two published *Drosophila* brain single-cell RNAseq datasets (Brunet Avalos et al., 2019; Ravenscroft et al., 2020). Our analyses reveal highly enriched expression of Obp44a in astrocytes, cortex glia, and ensheathing glia in first (Fig. 1E and fig. S2A) and third instar larval brains (fig. S2B). These findings highlight the abundant expression of Obp44a in the CNS glia, suggesting specialized roles beyond the traditional functions of odorant binding proteins (Larter et al., 2016; Rihani et al., 2021; Xiao et al., 2019).

To validate the findings of transcriptome analysis, we studied the distribution and dynamics of Obp44a protein in the fly brain. Obp44a enhancer Gal4 driven fluorescent markers targeting cell surface or nuclei revealed a broad distribution of Obp44a in the larval brain (Fig. 1F). Furthermore, we specifically labeled a subset of CNS glia, as indicated by co-staining with an antibody against Repo, a marker that labels glial nuclei (Fig. S3A). Conversely, immunohistochemical studies using a custom-made Obp44a antibody revealed a diffused distribution of the protein, which infiltrates the entire neuropil region in the larval brain (Fig. 1G). Additionally, high resolution imaging has revealed puncta labeled by the antibody, indicative of vesicles and secretory granules containing Obp44a.

Using CRISPR-mediated genome editing, we generated Obp44a::GFP knock-in lines to endogenously tag the Obp44a protein and visualize its *in vivo* localization (fig. S4). The Obp44a::GFP signal matches the antibody staining results closely, displaying a broad and diffused distribution within the larval brain neuropil region (Fig. 1H). Notably, although produced in astrocytes, Obp44a protein is low in astrocytes’ soma and high in the surrounding neuropil region (Fig.1H’), suggesting its effective release as a secretory protein. Similarly, in the adult brain, the Obp44a::GFP signal is found throughout the brain (Fig. 1I, left), with the visibly higher levels at the regions with densely packed neural fibers, such as the boundaries of antenna lobe (AL), antennal mechanosensory and motor center (AMMC), and subesophageal zone (SEZ). Furthermore, thin optic sections in the optic lobe region show the cytoplasmic distribution of Obp44a in the optic lobe chiasm (OLC) glia (Fig. 1I, right).

To verify the secretion of Obp44a from glia, we utilized Obp44a-Gal4 driven Nsf2 RNAi to specifically inhibit secretion from Obp44a expressing glia (Babcock et al., 2004; Coutinho-Budd et al., 2017) and examined the distribution of Obp44a::GFP. Indeed, blocking glia secretion resulted in altered Obp44a::GFP neuropile distribution, confining the protein within glia somas (Fig. 1J), thereby supporting the result that it is actively released from the glia. Furthermore, we conducted Nsf2 knock-down experiments in wildtype flies and detected the endogenous Obp44a protein using antibody staining. This revealed similar accumulation of Obp44a proteins in glia somas (fig. S3B), thus corroborating our observations of the knock-in lines and suggesting that the GFP tag does not significantly alter the secretory properties of the Obp44a protein.

### Obp44a is a fatty acid binding protein regulating brain lipid homeostasis

While homologues of Obp44a are exclusively insect odorant binding proteins (Rihani et al., 2021), the distribution of OBP44a suggests non-sensory roles. As one of the most abundant proteins produced by the CNS glia, what function does Obp44a serve? To answer this question, we first performed a protein structure homology search and identified *Aedes aegypti* (yellow fever mosquito) OBP22 (AeOBP22) as the closest structural homolog of Obp44a. Despite a mere 42% identity between their amino acid sequences, the predicated 3D structure of Obp44a closely resembles that of AeOBP22, featuring the six α-helices and a hydrophobic pocket region critical for ligand binding (Fig. 2A). Previous X-ray crystallographic and Nuclear Magnetic Resonance (NMR) studies have indicated the role of AeOBP22 as a fatty acid binding protein (Jones et al., 2019; Wang et al., 2020), thus prompting us to test the possible interactions between lipids and Obp44a.

**Figure 2.**
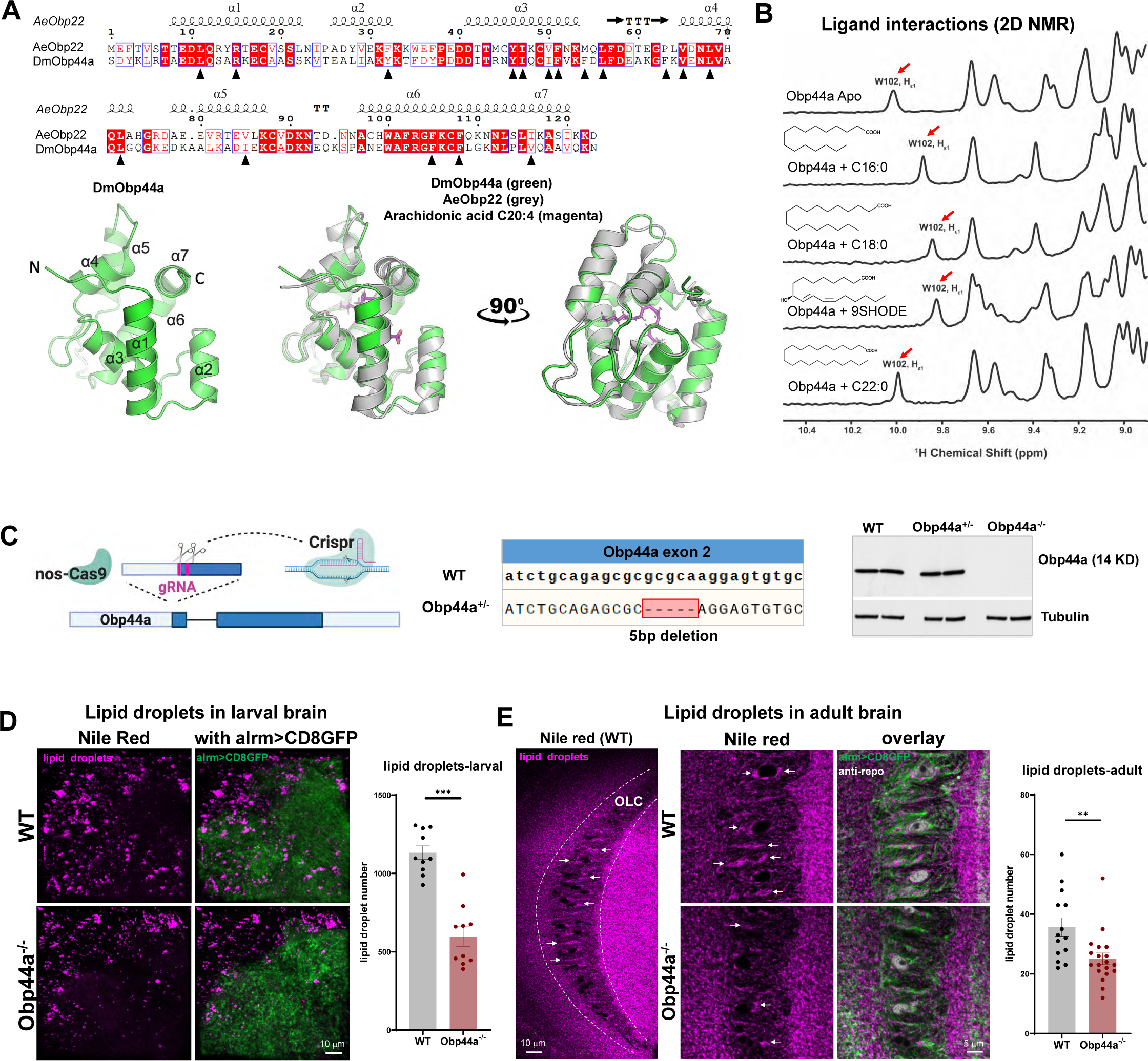
Obp44a is a fatty acid binding protein regulating lipid storage in the *Drosophila* CNS. (A) Protein structure prediction and homology analysis reveal similarities between *Drosophila* melanogaster Obp44a (DmObp44a) and Aedes aegypti Obp22 (AeObp22). Top: Protein sequence alignment without the signal peptides, highlighting critical amino acids (labeled with black triangles) forming the hydrophobic pocket in DmObp44a corresponding to the AAs identified in AeObp22. Bottom: AlphaFold2 predicted 3D protein structure reveals similarities between DmObp44a (green) and AeObp22 (grey), both featuring six alpha helices and a hydrophobic pocket that binds to fatty acid ligands, such as C20:4 arachidonic acid (magenta) shown in the model. (B) Fatty acid binding induced conformational changes in OBP44a are detected by NMR. The Trp102 side chain H_ε1_ proton (red arrow) is sensitive to the fatty acid ligand interactions. Distinctive shifts from the apo form (top panel) are detected in the presence of palmitic acid (C16:0), stearic acid (C18:0), and 9SHODE (C18:2), but not in the case of docosanoic acid (C22:0). (C) Generation of an Obp44a null mutant using CRISPR-Cas9 mediated gene editing (left) (created with BioRender.com), which produces a 5bp deletion in the second coding exon of Obp44a (middle). Western blots confirm the absence of Obp44a protein in the homozygous mutant (right). (D-E) Obp44a mutants have reduced numbers of lipid droplets in both the larval (D) and adult fly brain (E), detected by the Nile red staining (magenta). A significant reduction in lipid droplet numbers was found in Obp44a mutants at both stages. Astrocytes labeled using alrm>CD8GFP (green) are used as landmarks. Statistical significance assessed by unpaired t-test with Welch’s correction. **P<0.01, ***P<0.001. Error bars represent mean ± SEM; n=10 in (D), n=14,19 in (E).

To evaluate the lipid binding ability of Obp44a, we obtained a large amount of recombinant Obp44a protein and subjected it to NMR analysis, where conformational changes induced by ligand-protein interactions can be detected through the amide proton resonance repositions (He et al., 2023). For Obp44a, when a putative ligand interacts with its hydrophobic pocket, the bound state is characterized by the chemical shift of H_ε1_ NMR signal of Trp102 (W102) (Figure 2B). In the presence of fatty acids, including palmitic acid (C16:0) and stearic acid (C18:0), as well as the oxidized fatty acid 9(S)-HODE (C18:2), the H_ε1_ Trp102 (W102) signal consistently experiences an upfield shift (i.e. towards lower ppm values) from 10 ppm, corroborating a binding event (Figure 2B). However, the introduction of docosanoic acid (C22:0) elicits no such response, with the amide proton spectrum remaining the same as the one in the unbound apo state (Figure 2B). These results are consistent with previous findings in AeObp22, indicating that the hydrophobic pocket of Obp44a can accommodate a fatty acid ligand, in both native and oxidized forms, with an acyl chain length limited to 20 carbons.

Additionally, we conducted native gel shifting experiments using a BODIPY-labeled fluorescent C16-fatty acid (fig. S5). The incubation of C16-fatty acid and Obp44a resulted in the strong shifting of the fluorescent signal, providing additional evidence for interactions between Obp44a and fatty acids. It is worth noting that the major fatty acids present in the *Drosophila* brain range from C14 to C20 (Stark et al., 1993). The capability of Obp44a to bind these FA species supports the notion that it could play a role in regulating FA trafficking and metabolism within the fly brain. In addition, our data demonstrate the interaction between oxidized fatty acid 9(S)-HODE (C18:2) and Obp44a, suggesting a possible protective function for Obp44a against toxic lipids formed during oxidative damage in the brain (Pecorelli et al., 2019; Shen et al., 2021).

After establishing Obp44a as a protein that binds to intact and oxidized fatty acids, we sought to evaluate its potential function in fatty acid trafficking and storage *in vivo*. We generated a null mutant of Obp44a using CRISPR-Cas9 mediated gene editing (Fig. 2C) and examined the number of lipid droplets (LDs) in the mutant fly brain. The fly LDs, primarily consisting of triacylglycerols (TAGs) and sterol esters, serve as a major source of fatty acid storage and protect against oxidative damage during larval development (Kis et al., 2015; Papsdorf et al., 2023; Welte, 2015). Using the Nile red staining (Yin et al., 2021), we detected a 47% of decrease in the numbers of lipid droplets in the larval brain of Obp44a knockout flies (Fig. 2D). Similarly, in adult brains, quantifications of LD performed in the optic lobe chiasm (OLC) glia revealed a 30% of reduction of LD numbers in Obp44a mutants (Fig. 2E). Together, data obtained from larval and adult stages consistently support the function of Obp44a in maintaining the number of LDs in the CNS.

### Obp44a deficiency leads to morphological, physiological, and behavioral defects

Given the abundance and broad distribution of Obp44a in the fly CNS, as well as its affinity to the prominent species of fatty acids in the *Drosophila* system, we went on to determine the physiological consequence of its depletion at the cellular and behavioral levels. While Obp44a mutants are viable and have no gross morphological deficits, a close examination of the adult OLC glia layer revealed notable disruptions of the anatomical organization of glia (Fig. 3A). The OLC glia in wild-type flies displays evenly sized soma, minimal vacuolation, and a neatly aligned layer of nuclei, labeled by anti-Repo staining (Fig. 3A, top). In contrast, in Obp44a mutant flies, the OLC glia exhibited altered morphology characterized by irregular soma shapes, the presence of large vacuoles (Fig. 3A, arrows) and disorganized nuclei (Fig. 3A, bottom, stars).

**Figure 3.**
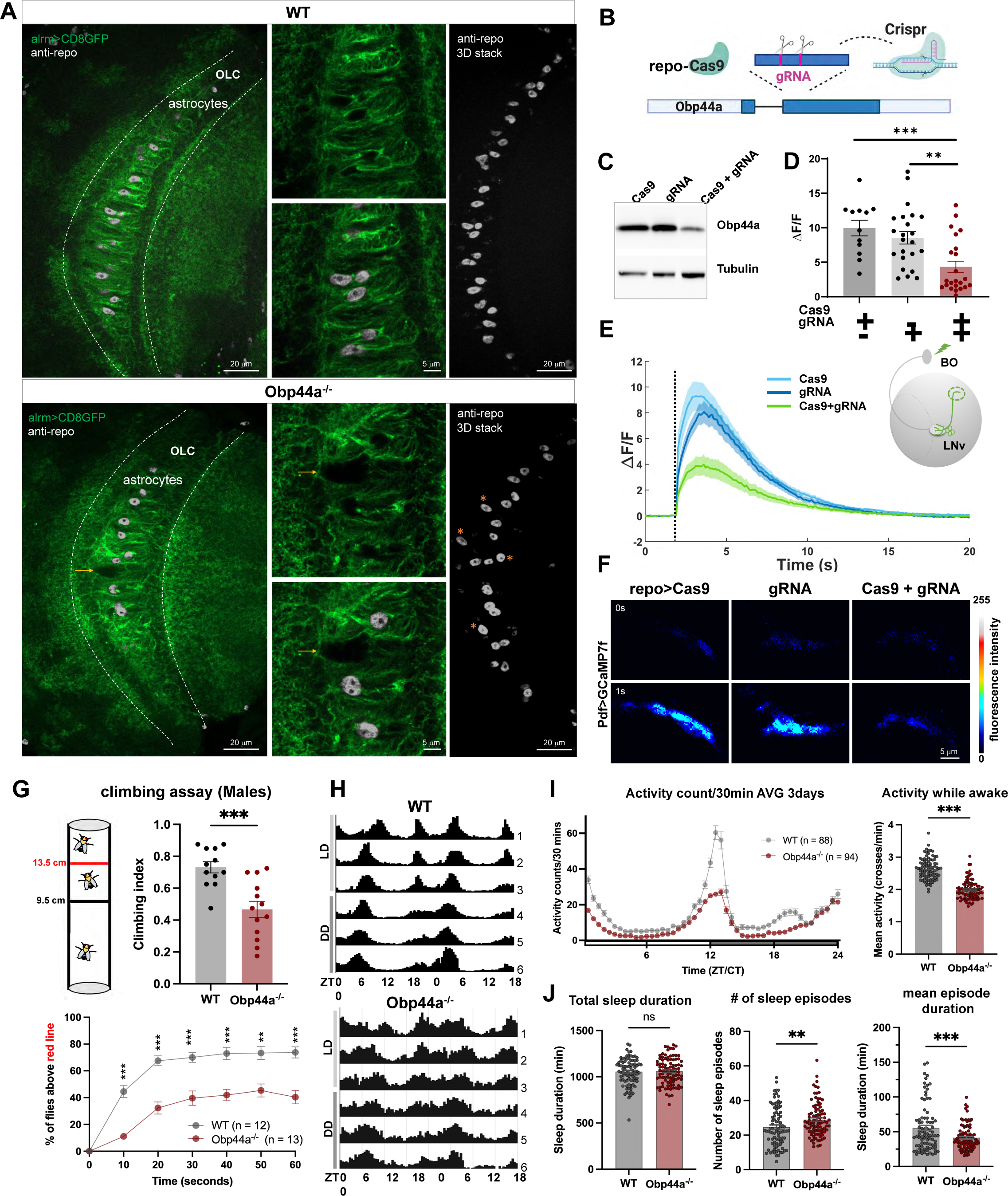
Morphological, physiological, and behavioral deficits in Obp44a mutants. (A) Obp44a mutants show disrupted morphology of astrocytes (green channel) and the arrangement of astrocyte nuclei (grey channel) in the OLC region adult brains. Presence of large vacuoles and disorganized nuclei in Obp44a mutant are marked by orange arrows and orange stars. (B-C) CRISPR/Cas9-mediated tissue-specific mutagenesis reduces Obp44a levels in glia. Schematic diagram (B) and western blot confirmation of reduced Obp44a levels (C). (D-F) Glia-specific Obp44a mutagenesis reduces light-elicited calcium responses in LNvs, suggesting a role of Obp44a in supporting neuronal physiological responses. (D) Quantification of peak changes in GCaMP signal induced by light stimulation (ΔF/F). Statistical analysis performed using one-way ANOVA followed by Tukey post-hoc test. **P<0.01, ***P<0.001. Error bars represent mean ± SEM; n=12-23. (E) Traces depicting average GCaMP signals recorded at the axonal terminal region in LNvs. The shaded area represents SEM, and the dashed black line indicates the 100 ms light pulse used for stimulation. Right panel: The schematic diagram illustrating the setup for calcium imaging experiments in LNvs. The light pulse through the Bolwig’s organ (BO) elicits calcium responses in LNvs, which are recorded at the axonal terminal region (dashed green circle). (F) Representative frames of Pdf>GCaMP7f recordings at 0 and 1 second after stimulation for controls and glia-specific knockout. (G) Obp44a mutants exhibit significantly reduced climbing ability, indicated by a reduction in climbing index (top) and climbing height (bottom). Statistical significance was determined using an unpaired t-test with Welch’s correction for climbing index and Welch’s ANOVA test with Dunnett’s multiple post-tests for climbing curve, **P<0.01, ***P<0.001. Error bars represent mean ± SEM; n=12 and 13 groups with 20 flies per group. (H) Altered locomotion patterns for Obp44a mutants in light: dark (LD) and constant dark (DD) conditions. Representative actograms of average activity are shown. (I) Daily locomotor activity profiles and quantifications demonstrating reduced activity levels in Obp44a mutants. (J) Quantifications of total sleep duration, number of sleep episodes, and episode duration. Comparing to the wildtype controls, Obp44a mutant flies exhibit an increased number of sleep episodes and decreased episode durations. Statistical significance was assessed using an unpaired t-test with Welch’s correction. **P<0.01, ***P<0.001, “ns” indicates not significant. Error bars represent mean ± SEM; n=88, 94.

To probe whether Obp44a depletion influences neuronal function, we evaluated the physiological responses of larval ventral lateral neurons (LNvs), where the light activation of presynaptic photoreceptors leads to a large increase in calcium influx in the postsynaptic LNvs, reported by the calcium indicator GCaMP7f (Dana et al., 2019; Shemesh et al., 2020; Yuan et al., 2011). Using a combination of Repo enhancer driven Cas9 expression (Repo>Cas9) and a gRNA transgene specifically targeting Obp44a, we achieved glia-specific knock-down of Obp44a and confirmed its efficiency by western blots (Fig. 3B, C). Obp44a knock-down led to a significant reduction of light induced physiological responses in LNvs, indicated by over 50% of reduced amplitude of calcium transients, as compared to both control groups (Fig. 3D-F), suggesting that Obp44a is required for normal neuronal physiological responses.

Next, to assess the impact of Obp44a deficiency at the organismal level, we conducted behavioral studies. Notably, mutant flies lacking Obp44a tend to stay at the bottom of the food vials, in contrast to wild-type control flies which typically climb to the top of the vials. Quantifications of our climbing assay confirmed this observation and demonstrated that Obp44a mutants have reduced ability to climb and perform negative geotaxis (Fig. 3G and fig. S6). Furthermore, to gain additional information regarding their locomotion, circadian rhythm, and sleep behavior, we tested Obp44a mutants using the *Drosophila* Activity Monitor (DAM) system. Consistent with the findings of the climbing assay, Obp44a knockout flies showed significantly reduced locomotor activity during their active periods (Fig. 3H, I). In addition, although their circadian rhythm appears normal (Fig. 3H), the mutant flies exhibited fragmented sleep patterns (Fig. 3J). While their total sleep time is similar to the control flies, Obp44a knockout flies have increased number of sleep episodes and shorter sleep episode duration, suggesting a function of Obp44a in maintaining normal locomotor activities and facilitating sleep consolidation.

Overall, our anatomical, physiological, and behavioral analyses strongly support Obp44a as an integral part of the molecular machinery required for normal structure and function of the fly brain and the general fitness of the animal. Notably, the lipid storage phenotype, the physiological and behavioral deficits in Obp44a mutants are analogous to the ones found in FABP7 knockout mice (Gerstner et al., 2017; Owada et al., 2006; Shimamoto et al., 2014), suggesting *Drosophila* Obp44a and mammalian FABP7 are likely functional homologues.

### Obp44a regulates lipid storage and redox homeostasis in the fly brain

FABPs facilitate intra and inter cellular lipid trafficking in both vertebrate and invertebrate systems (D’Anneo et al., 2020; Kis et al., 2015). Given the intricate relationship among different metabolic pathways and complex expression patterns of distinctive types of FABPs, their impact on overall metabolism and brain lipid homeostasis remains unclear. Since Obp44a is abundant and has expression exclusive to the fly CNS, we took this opportunity to address these questions experimentally and investigated metabolomic changes associated with the Obp44a deficiency.

Using Hydrophilic Interaction Liquid Chromatography Mass Spectrometry (HILIC-MS) analysis (Contrepois et al., 2015) of larval brains (Fig. 4A, B), we detected alterations in ∼300 metabolites in the Obp44a mutant samples. Among these, 141 metabolites exhibited increased levels, while 159 showed reduced levels compared to the wild-type controls (table S5). These changed metabolites span a broad range of molecular categories (Fig. 4C), indicating widespread metabolic perturbations in the absence of Obp44a. Notably, among the top 28 metabolites displaying a significant change (p<0.05) across biological replicates, several groups of molecules stood out, including upregulated 13-HODE and Phosphatidylethanolamines (PEs), alongside downregulated phosphatidylinositols (PIs), diacylglycerols (DAGs) and carnitines (Fig. 4C, D; fig. S7B). The downregulated lipid categories bear particular significance as they are directly linked to Obp44a’s fatty acid binding capability, consistent with the critical roles of fatty acids in plasma membrane turnover and signaling processes. Furthermore, a notable reduction in glycogen levels is also noted, suggesting a shift in the brain energy supply within the mutant animals.

**Figure 4.**
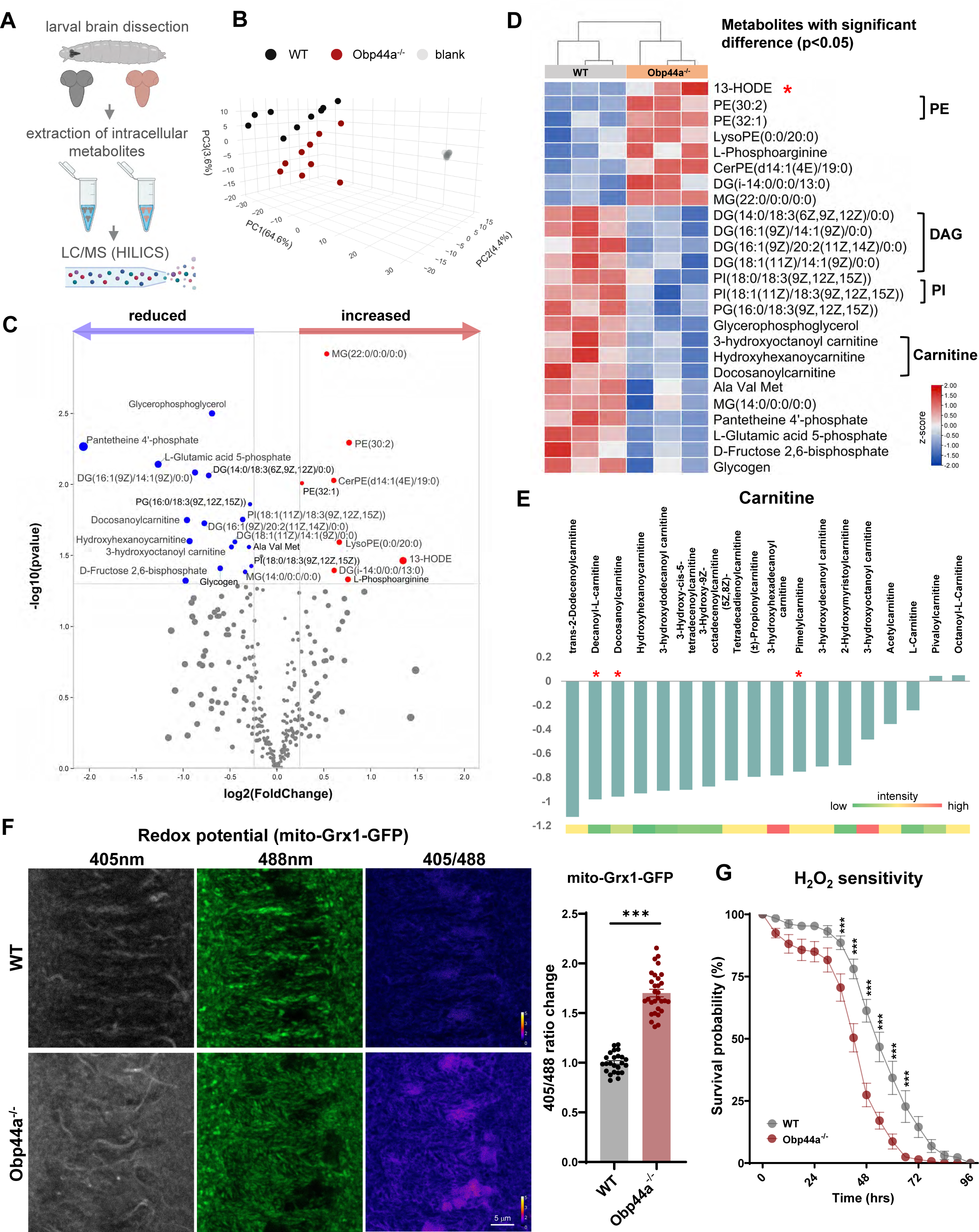
Obp44a regulates brain lipid and redox homeostasis. (A) Schematic diagram illustrating the sample preparation procedure of larval brain metabolomics analysis through hydrophilic interaction liquid chromatography (HILIC) mass spectrometry. (B) Principal Component Analysis (PCA) of the larval brain metabolome across all biological and technical replicate samples for wildtype, Obp44a mutant and blank controls. (C) Volcano plot illustrates total detected metabolite changes in WT and Obp44a^-/-^ larval brain. Red and blue dots represent increased or decreased metabolites meeting the p-value cutoff (p-value<0.05) and log2 fold change (LFC>0.25) criteria. (D) Heatmap displays metabolites with significant changes in Obp44a^-/-^ mutant brain, including PE, DAG, PI, carnitines, and oxidated fatty acid 13-HODE. (E) Consistent reduction of multiple species of carnitines were detected in the Obp44a mutant brains. Red stars indicate p-value<0.05. The color-coded bar (bottom) illustrates the average intensity of each metabolite within this category. (F) Glutathione redox (GSH/GSSG) biosensor (mito-roGFP2-Grx1) analysis reveals elevated redox potentials in the Obp44a mutant brains. Representative raw and ratiometric images of adult optic lobes expressing mito-roGFP2-Grx1 are shown (left). Quantification of 405/488 ratiometric fluorescent changes in the optic lobe region indicates an elevated redox potential in the mutant brain. Statistical significance is assessed by unpaired t-test with Welch’s correction. ***P<0.001. Error bars represent mean ± SEM; n=24, 30. (G) Survival curve of adult WT and Obp44a^-/-^ flies subjected to 5% H_2_O_2_ treatment. Obp44a^-/-^ flies exhibit significantly reduced survival probability after 24-hour treatment. Statistical significance assessed by Welch’s ANOVA test with Dunnett’s multiple post-tests, ***P<0.001. Error bars represent mean ± SEM; n=9 groups with 16 flies per group.

In particular, 13-hydroxyoctadecadienoic acid (13-HODE), a major lipoxygenation product synthesized from linoleic acid (Pecorelli et al., 2019; Shen et al., 2021), emerges as one of the most significantly elevated metabolites in Obp44a mutants (Fig. 4C, D), indicating an increased level of oxidized lipids associated with the loss of Obp44a. Additionally, Obp44a mutants exhibit a significant downregulation of four DAGs (Fig. 4C, D), which serve as crucial phospholipid precursors and components of cell membranes (Eichmann & Lass, 2015), as well as intermediates for TAG (fig. S7A), the primary form for fatty acid storage and a major component of lipid droplets (Barbosa et al., 2015; Fujimoto & Parton, 2011). Upon examining all TAGs and DAGs detected in the metabolomics data, we consistently observed a reduction of these lipids in Obp44a mutants, with only a few exceptions (fig. S7A). In combination with the reduced lipid droplet phenotype that we observed in Obp44a mutants (Fig. 2D, E), these data suggest a reduced lipid storage caused by Obp44a deficiency in the fly brain.

Another notable group showing significant and consistent changes in Obp44a mutants are carnitines. Out of the eighteen carnitines detected in the metabolomics dataset, sixteen of them, including L-carnitine, exhibited reduction and two remained unchanged (Fig. 4E). L-carnitine serves as a cofactor for the transport of long-chain fatty acids (i.e. more than 14 carbons) into the mitochondria for subsequent β-oxidation and energy production (Longo et al., 2016). The depletion of carnitine in Obp44a mutants suggests a deficit in intracellular long chain fatty acid transport and increased levels of β-oxidation, possibly from peroxisome activity. Indeed, the high levels of 13-HODE are consistent with peroxisomal 15-LOX-1 activity (Jo et al., 2020).

The metabolomics analysis indicates that the deficiency of Obp44a leads to diminished lipid storage, altered lipid transport for metabolism within the mitochondria, and elevated lipid oxidation. To investigate the involvement of Obp44a in regulating the oxidative state of the brain, we performed several experiments. First, we evaluated the redox potential in Obp44a mutant flies utilizing an in vivo redox biosensor mito-Grx1-roGFP under the control of a tubulin enhancer (Albrecht et al., 2011). Remarkably, we observed a significantly higher fluorescence ratio change (405/488) in the Obp44a mutant fly brains, indicating an elevated glutathione redox potential (EGSH) in the mitochondria, which are major sources of oxidants in cells (Fig. 4F), consistent with a heightened oxidative state in the mutant brain. In addition, we conducted live imaging experiments on larval brains, measuring superoxide level in astrocytes using the fluorescent probe Dihydroethidium (DHE) (Bailey et al., 2015). As anticipated, we detected an elevated fluorescent intensity in the astrocyte nuclei of Obp44a mutant flies compared to wild-type controls (fig. S8). Furthermore, to assess the flies’ sensitivity to oxidative stress, we subject them to H_2_O_2_-induced oxidative stress via feeding, which typically results in lethality in wildtype flies within approximately four days. In comparison, Obp44a mutants displayed increased sensitivity to H_2_O_2_ feeding, with significantly reduced median survival times (Fig. 4G). This outcome suggests that elevated oxidative stress in the Obp44a mutant renders the animal more susceptible to H_2_O_2_ challenge. Collectively, our data provide evidence supporting the role of Obp44a in maintaining redox homeostasis and reducing oxidative stress in the *Drosophila* brain.

### Obp44a mediates lipid efflux in the developing fly brain

During the larval and pupal development, the *Drosophila* CNS undergoes two waves of neurogenesis, accompanied by significant brain remodeling, mass synaptic pruning, and programmed cell death associated with metamorphosis. These essential developmental processes produce abundant cellular debris, with lipids being a major component. Given Obp44a’s affinity for fatty acids, and its high expression in the larval brain, we hypothesize that it may play a role in mediating lipid recycling and efflux. To test this hypothesis, we utilized the Obp44a::GFP knock-in line to observe the trafficking of the Obp44a protein and its fatty acid cargo in an *in vivo* setting.

As shown previously, endogenously tagged Obp44a::GFP exhibits a diffused distribution throughout the entire neuropil region. This pattern can be replicated by expressing a GFP tagged Obp44a in either neuron or glia in the larval brain using cell-type specific enhancers. The results demonstrate Obp44a’s ability to be effectively secreted and enter both cell types (Fig. 5A). Using acutely dissociated brain cells, we confirmed this observation by finding glial-derived Obp44a::GFP in neurons and *vice versa* (Fig. 5 B, C).

**Figure 5.**
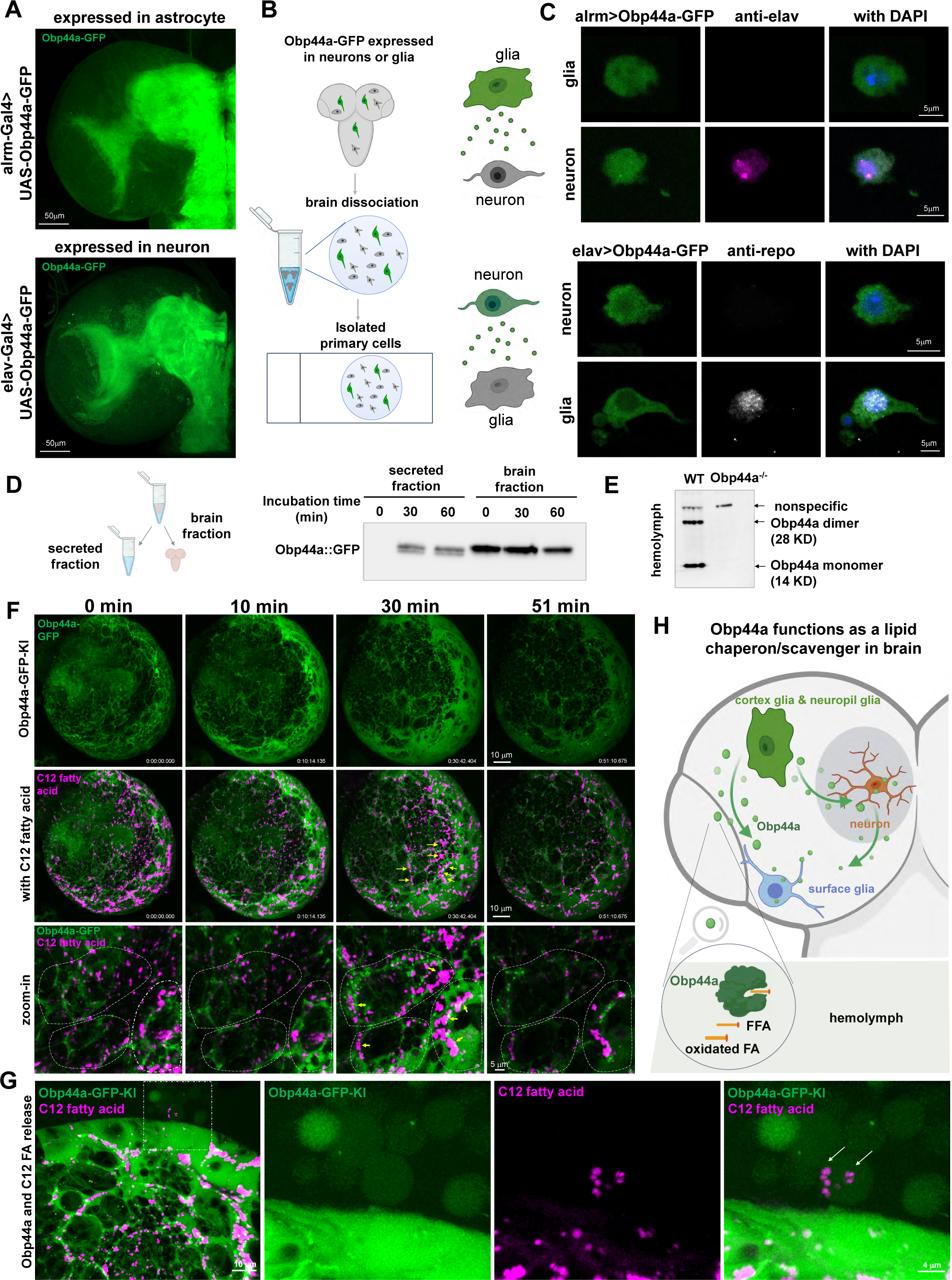
Obp44a mediates brain fatty acid trafficking and efflux. (A) Expression of Obp44a::GFP in astrocytes or neurons recapitulates the protein localization observed in the Obp44a::GFP knock-in line, demonstrating efficient secretion and dynamic trafficking of Obp44a across different cell types. (B-C) Obp44a traffics between glia and neuron. (B) Schematic illustration of experiments using acutely dissociated brain cells to visualize the distribution of Obp44a::GFP specifically produced in glia or neuron. (C) Obp44a::GFP produced in astrocytes (alrm-Gal4>UAS-Obp44a::GFP) is observed in elav-positive neurons, while neuronal expressed Obp44a::GFP (elav-Gal4>UAS-Obp44a::GFP) is detected in repo-positive glia, indicating Obp44a’s ability to traffic between glia and neuron *in vivo*. (D) Schematic diagram (left) illustrating the western blot analysis (right) performed on larval brain explants and the medium collected after incubating explants for 30 to 60 min. The result demonstrates that Obp44a::GFP is secreted outside of the brain into the culture medium. (E) Monomeric and dimeric Obp44a proteins are detected in the larval hemolymph. (F) Time-lapse live imaging of the larval brain explant reveals the mobilization of fatty acids accompanied by Obp44a::GFP migration from the brain neuropil region surface glia. Top two panels: representative MIP images of a 3^rd^ instar larval brain from the Obp44a::GFP knock-in line are show. Fluorescently labeled C12 fatty acid signal (magenta) is distributed across the brain region at the beginning of the imaging session and starts to mobilize toward the surface glia, together with the changes of Obp44a::GFP localization from the central to the surface brain region. The arrows indicate the C12-fatty acid accumulation in surface glia at 30min, which is diminished at 51 min. Bottom panel: Zoom-in single optic sections illustrate the mobilization of both C12-fatty acid and Obp44a::GFP into three surface glia (with dash outlines). (G) Obp44a facilitates fatty acid efflux through secretion into the hemolymph. 30 min after imaging session started, Obp44a::GFP is secreted outside out the brain via exosomes, which also contain labeled C12-fatty acid (magenta). (H) Schematic diagram illustrating Obp44a mediating fatty acid trafficking between neuron and glia and facilitate lipid efflux through secretion into the hemolymph. Produced in cortex glia and neuropil glia, Obp44a is effectively secreted from glia into the neuropil region and traffic through various brain cells, such as neurons and surface glia, eventually being released into the hemolymph. Acting as a lipid chaperon and scavenger, the abundant glial production of Obp44a contributes to the regulation of fatty acid trafficking and metabolism, as well as to the clearance and efflux of oxidized fatty acids, both are critical for a healthy brain environment.

Next, we conducted experiments to examine whether Obp44a traffics outside of the brain. Incubating the larval brain in physiological saline for 30-60 minutes resulted in the ready detection of Obp44a::GFP in the supernatant (Fig. 5D). Moreover, by performing western blots in hemolymph collected from the larvae, we found Obp44a in both dimeric and monomeric forms in wildtype animals, but not in the null mutants (Fig. 5E). Because only the monomeric form of the Obp44a protein is detected in the larval brain extract, this result suggests that Obp44a monomer is likely the functional form of protein, while its oligomers are utilized for trafficking within the hemolymph. In addition, we observed strong Obp44a::GFP signals in the nephrocytes, the “kidney-like” cells on the larval epidermis that are involved in filtering the hemolymph, supporting Obp44a’s presence in the circulatory fluid (fig. S4).

To examine how Obp44a mobilizes fatty acid cargos *in vivo*, we performed live imaging experiments using larval brain explants. In these experiments, we incorporated a fluorescently labeled C12 fatty acid (C12-red) along with GFP-tagged Obp44a. After 24 hours of feeding, C12-red was readily detected in brain tissue, primarily in droplet-like structures surrounded by Obp44a::GFP signals. Remarkably, upon dissection, Obp44a::GFP signals in the brain explants rapidly diminished from the neuropil region and were transported outward to surface glia (Fig.5F, top), where they appeared to be secreted into the hemolymph through exosome like vesicles (Fig. 5G, green channel). The C12-red signal exhibited a similar trend of movement, migrating from central brain regions towards surface glia and eventually disappearing (Fig.5F, middle and bottom, fig. S9). Interestingly, the C12-red signal was observed to exit the brain within vesicles filled with Obp44a::GFP, demonstrating the efflux of fatty acids accompanied by Obp44a mobilization (Fig. 5G).

The rapid mobilization of Obp44a::GFP prompted us to investigate its response to oxidative stress in intact animals. To induce oxidative stress in adult flies, we subjected Obp44a::GFP knock-in flies to a strong oxidant tert-butyl hydroperoxide (tBH) (Obata et al., 2018), which lead to lethality in adult flies within two days. Intriguingly, upon 24 hours of feeding with tBH, Obp44a::GFP distribution is drastically altered, resulting in the accumulation of numerous droplet-like structures containing Obp44a::GFP in various intracellular regions of the fly brain (fig. S10). Upon removal of tBH from the food, most of the cumulated Obp44a::GFP signals dispersed after four days (fig. S10). These data suggest that Obp44a act as a lipid scavenger to collect oxidized fatty acids and facilitate their outward transport into the hemolymph.

Our integration of endogenous labeling and live imaging experiments have revealed the dynamic trafficking of Obp44a among diverse cell types within the brain. Obp44a is initially secreted from cortex and neuropil glia into the extracellular space, then infiltrates neurons and neuropil regions, facilitating the transport of fatty acid and protecting them from oxidative damage. This process supports crucial functions such as membrane biogenesis, energy production, and lipid storage. Under conditions of oxidative stress, Obp44a effectively sequesters oxidized lipids, exiting the brain through surface glia and hemolymph. With its affinity for both native and oxidized fatty acids, coupled with its abundant production in glial cells and rapid mobilization, Obp44a plays a pivotal role in regulating intracellular fatty acid levels and trafficking. Furthermore, its action likely contributes to a sustained lipid efflux in the brain, crucial for the elimination and recycling of oxidized lipids and membrane components from cellular debris, thereby ensuring the establishment of a healthy lipid environment during brain development (Fig. 5H).

## Discussion

Fatty acids play crucial roles in structural, signaling and metabolic processes (Chung et al., 2023; Dutta et al., 2023). Within the CNS, they are closely associated with a spectrum of physiological and pathological conditions (Falomir-Lockhart et al., 2019; Wang et al., 2022). As primary carriers facilitating the transport of hydrophobic fatty acids, FABPs have undergone extensive investigations through in vitro analyses, revealing their 3D structures, ligand binding affinities, and interactions with phospholipid containing vesicles and membranes (Storch & Corsico, 2023). Furthermore, their specific distributions across tissues and cell types, along with phenotypic analyses of knockout animals, suggest their essential roles in supporting brain function (Asaro et al., 2021; Cheng et al., 2021; Killoy et al., 2020). However, what has been lacking is the *in vivo* assessment of how FABPs impact brain metabolism, interact with lipid cargos, and respond to cellular environments and membrane structures. In this study, we identified Drosophila Obp44a as a highly expressed glia derived secretory FABP and began to address these questions. Specifically, we observed that Obp44a deficiency led to widespread changes in the larval brain metabolome. On one hand, consistent reductions in TAG and DAG levels in mutants indicate a general decrease in brain lipid storage, likely resulting from disrupted fatty acid trafficking. On the other hand, the depletion of carnitines and alterations of diverse species such as DAG, PI and carnitines suggest substantial impacts on mitochondrial metabolic β-oxidation, membrane composition, and signal transduction cascades. These changes have multifaceted physiological consequences that could be further investigated through additional molecular and biochemical studies.

Surprisingly, we identified Obp44a as having the functional characteristics of FABPs but without their structural similarity. Indeed, multiple members of FABPs have been subjected to X-ray crystallography, NMR, and other biochemical and biophysical studies. Despite significant divergence in their protein sequences, all known FABPs, including the dFABP, share nearly identical 3D structures characterized by a 10 stranded antiparallel β-barrel (Storch & Corsico, 2023). The binding pocket for fatty acids is located inside this β-barrel, with the N-terminal ‘cap’ domain positioned at the opening (Furuhashi & Hotamisligil, 2008). In contrast, the *Drosophila* Obp44a protein shares structural features similar to other odorant binding proteins from insect, containing an internal hydrophobic cavity formed by six alpha-helices (Larter et al., 2016). Moreover, variations in the C-terminal tails of insect Obps affect their abilities to bind and release ligands, ranging from small organic molecules to fatty acids. For example, the flexible C-terminal region of AeObp22 creates an expandable binding pocket that accommodates long-chain fatty acids with a carbon number over 12 (Wang et al., 2020). Although the high-resolution 3D structure of Obp44a remains unsolved experimentally, structural alignment and NMR studies clearly indicate its capacity for fatty acid binding, akin to AeObp22. These structural distinctions from classical FABPs prompt us to ask questions about the diversity in 3D structures of proteins that bind to fatty acids. Not only do proteins with similar 3D structures bind to distinct ligands, as evidenced by insect Obp family members, but proteins with different 3D structures also can bind to the same ligand, such as the case for AeObp22, DmObp44a, and mammalian FABPs. Ultimately, the discovery of new lipid binding proteins will likely rely on a combination of sequence and structure homology searches with unbiased systematic biochemical and functional screens. This synergistic approach could potentially lead to a substantial expansion of our existing knowledge of lipid-protein interactions. In particular, the structural and functional differences between OBP44a and previously characterized FABPs suggest that the unique functions of OBP44a such as its secretory properties could be supported by its α-helical conformation.

Our research has uncovered an innovative strategy employed by the invertebrate system to regulate the lipid environment within the developing CNS. When we compared the expression level and tissue distribution of Obp44a with the dFABP and three fatty acid transfer proteins found in the *Drosophila* genome, dfatp1-3 (fig. S11), we confirmed that Obp44a is specifically expressed in the brain and showed that it is the dominant FABP in the brain, particularly during the larval stage. Furthermore, a key characteristic distinguishing Obp44a from other FABPs is the presence of a N-terminal signal peptide, facilitating its efficient secretion from both glia and neurons, and allowing Obp44a to have an extracellular role as well. While dFABP is also produced in cortex and surface glia, it is primarily localized in the glia soma, not in neuron or neuropil regions, indicating its specific role in regulating intracellular fatty acid transport and lipid droplet formation within glia (Kis et al., 2015). Additionally, fatty acid transport proteins (FATPs) are transmembrane proteins involved in the uptake of long-chain fatty acids from extracellular fluid (D’Anneo et al., 2020; Mitrofanova et al., 2021; Pohl et al., 2004). Consequently, the abundance and secretory nature of Obp44a enables the protein to function as an efficient chaperone for both inter- and intracellular lipid transport. Moreover, it acts as a scavenger for capturing and clearing toxic lipids in the larval brain, where significant tissue growth, neurogenesis, and synaptic remodeling occur rapidly over a few days, creating a high demand for stabilizing factors that maintain lipid and redox homeostasis. These functions are likely fulfilled by Obp44a collaborating with other FABPs during this critical developmental period. Nevertheless, the anatomical, physiological, and behavioral deficits associated with Obp44a deficiency and its broad impact on the larval brain metabolome indicate that Obp44a is a major contributor to brain health during development and under oxidative stress conditions.

The utilization of Obp44a as a CNS-specific FABP stands as a great example of naturally occurring protein engineering within the evolution trajectory of the *Drosophila* genome. Whether other animal species have evolved comparable or divergent strategies to address challenges associated with fatty acid trafficking under either physiological or pathological conditions poses interesting questions for future studies. Notably, both insect and mammalian Obps have been exploited for biotechnological applications. Their capability to interact with a wide array of hydrophobic molecules, coupled with their small size, stability, and solubility, renders Obps suitable targets for various applications, such as developing insect repellent and biosensors (Brito et al., 2020). Although the cell biological process involved in the secretion and trafficking of Obp44a remains to be determined, a tremendous potential exists to leverage its unique properties to regulate lipids in specific disease models or engineer novel drug delivery vehicles for the nervous system.

## ACKNOWLEDGMENTS

We thank Dr. Lucy Forrest at NINDS for facilitating the protein structure homology search; and Dr. Chun Han at Cornell University for helpful discussions; and Dr. Carolyn Smith at the NINDS Light Imaging Core, Dr. Sarah K. Williams Systems at NIMH Imaging Resource Core, Dr. Ling Yi and Dr. Vincent Schram at NICHD Microscopy and Imaging Core, Kelly Veerasammy and Malia Morioka at CUNY ASRC for technical support. Dr. Rinat Abzalimov is supported by PSC-CUNY Research Award Program. Dr. Ye He and Lele Xu are supported by NIH R01DK136013 and NIH R01NS12654. Dr. Qiangjun Zhou is supported by NIH R00 MH113764 and R01 MH132918. This research is supported by the intramural research program of National Institutes of Health. Project number 1ZIANS003137.

## Materials and Methods

### Fly stocks

Fly stocks are maintained in the standard cornmeal-based fly food in a 25°C incubator with humidity control. Larvae and adults are cultured in the light: dark (LD) condition with a 12-hour light: 12-hour dark light schedule. Unless otherwise noted, all larvae were collected between ZT1-ZT3 (ZT: zeitgeber time in a 12:12 hr light dark cycle; lights-on at ZT0, lights-off at ZT12). The following Drosophila melanogaster stocks were used for experiments: Obp44a-gRNA, UAS-Obp44a::GFP, UAS-CD2::mCherry, Obp44a::GFP CRISPR knock-in, and Obp44a CRISPR knock-out, are generated in this study. Obp44a-Gal4 (GMR90C03; BDSC 47122); UAS-CD8::GFP (BDSC 5137); UAS-RedStinger (BDSC 8547); Canton-S (wild-type for this study; BDSC 64349); UAS-Nsf2-RNAi (BDSC 27685); alrm-Gal4 (gifted by Dr. Marc Freeman); repo-Gal4 (BDSC 7415); elav-Gal4 (BDSC 458); repo-Cas9 (gifted by Dr. Chun Han); Pdf-Gal4 (BDSC 6899); UAS-GCaMP7f (BDSC 86320); tub-mito-roGFP2-Grx1 (BDSC 67669)

### RNA-seq data Collection and analysis

The single-cell RNA-seq data at the first instar larval stages (Fig. 1A-B, 1E, S1C and S2A) were from a previous publication and can be accessed via the following link: (https://cells.ucsc.edu/?ds=dros-brain) (Brunet Avalos et al., 2019). Similarly, the third instar larval brain single-cell RNA-seq data (Fig. S2B) were obtained from (https://github.com/aertslab/SCopeLoomR) as detailed in the published work (Ravenscroft et al., 2020). The third instar (Fig. 1C, S1A and S1C) and adult astrocyte RNA-seq data (Fig. 1C, S1B and S1C) were retrieved from published report (Huang et al., 2015). RNA-seq data of the Drosophila antenna (Fig. 1D) was derived from the published study (Menuz et al., 2014). The third instar CNS data (Fig. 1D) were from ModEncode (ID-4658) [http://data.modencode.org/cgi-bin/findFiles.cgi?download=4658] and analyzed according to our previous study (Yin et al., 2018). RNA-seq data corresponding to adult female and male Drosophila brains (Fig. 1D) were retrieved from NCBI under the accession code GSE153165, as reported (Nandakumar et al., 2020). RNA-seq data (Fig. S11) analyzed for tissue-specific expression levels of Obp44a, Fabp, and Fatp, and the RPMK values were extracted from FlyBase, originating from various sources including NCBI and ModEncode (PRJNA75285, SRP003905, modENCODE_3207).

### Seurat data processing and secretome analysis

We used the Seurat version 4.0 pipeline (Hao et al., 2021) to process single-cell RNA-seq datasets from both first and third instar larval Drosophila brains. The first instar larval brain dataset (Fig 1A-C, E and fig S1C) underwent a previously published processing method (Brunet Avalos et al., 2019), involving filters to retain cells exhibiting unique feature counts ranging between 200 and 4500, while also restricting mitochondrial gene content to less than 20%. This filtration yielded 4349 cells, encompassing a total of 12,942 detected genes.

The third instar dataset (Fig 1C and fig S1A, C), conversely, had undergone an initial filtration step, incorporating several quality control criteria according to their respective publication (Ravenscroft et al., 2020). This rendered a dataset comprising 5056 cells, encompassing 9853 detected genes. Subsequent to the construction of Seurat objects, standard pre-processing steps were executed before delving into downstream analysis. This included log-normalization, employing a scale factor of 10,000, to standardize gene expression across individual cells by considering the total gene expression within each dataset. A subsequent linear transformation was applied. To determine highly variable genes, we implemented the FindVariable Features function with default parameters, following the guidelines provided by the R package developer.

To ascertain the true dimensionality of the dataset, we explored several strategies, including the Elbow-Plot and JackStraw-Plot tests, in conjunction with an evaluation of PC-heatmaps. Ultimately, we retained 31 and 35 dimensions for the first and third instar datasets, respectively, employing these dimensions for cell cluster identification through a graph-based approach. The resolution parameters of 3 and 2.5 were utilized when employing Umap for visualization in these two datasets.

For the analysis of the Drosophila secretome dataset, encompassing 1709 secreted proteins, we drew upon the FlyXCDB online database (Pei et al., 2018). In the first instar astrocyte cell cluster, comprising data from a prior study (Brunet Avalos et al., 2019), a total of 258 secreted proteins were identified (Table S1). Similarly, in the third instar larval astrocyte-specific dataset (Huang et al., 2015), 176 secreted proteins were discerned (Table S2). Finally, in the adult astrocyte-specific dataset (Huang et al., 2015), 29 secreted proteins were detected based on the original paper threshold of Log2FC change ≥ 0.5 (Table S3). These findings are visualized through heatmaps and dot plots in Fig 1A-C and Fig S1.

### Generation of Obp44a::GFP knock-in line

The Obp44a::GFP knock-in line was generated through the integration of a donor plasmid (pTEGM) (gifted by Dr. Fengqiu Diao and Dr. Benjamin White) (Diao et al., 2015) and a gRNA plasmid (gifted by Dr. Chun Han) (Poe et al., 2019), with the latter containing two gRNA sequences. The donor plasmid consisted of three crucial fragments: a synthesized DNA sequence encompassing the 560bp Obp44a 5’UTR sequence plus a 34bp FRT sequence, the Obp44a coding sequence containing the coding region (CDS) and its only intron obtained from genomic material, and a synthesized DNA sequence comprising a 34bp FRT sequence plus the 620bp Obp44a 3’ UTR sequence. The two PAM (Protospacer Adjacent Motif) sites utilized for Cas9 cleavage were rendered non-functional within the synthesized sequence. These synthesized 5’ UTR, Obp44a coding sequence and 3’ URT sequences were introduced for recombination with the endogenous Obp44a genomic sequence, thereby replacing the native Obp44a locus.

To generate the gRNA expression vector, we used a modified gRNA cloning vector based on pAC-attB-CaSpeR4 (Volkenhoff et al., 2015). This vector, (a gift from Dr. Chun Han) featured a U6:3 promoter and two gRNA-expression cassettes arranged in tandem (tRNA + gRNA + gRNA core EF) (Poe et al., 2019). The specific targeting sequences for Obp44a within the gRNA expression vector were “ccgagcgagcattcagtcctca” and “ccgctcaggctctgcaatcctac.” Both the donor and gRNA plasmids were prepared at 0.5 to 1 μg/μl concentration for co-injection into the nos-Cas9 (attP2) fly line (by Genetivision). The resulting knock-in line was validated through DNA sequencing and western blots.

### Generation of Obp44a::GFP knock-in line

The generation of the Obp44a mutant involved the use of the nos-Cas9 (attP2) fly line and a Obp44a-gRNA line in the published collection (Meltzer et al., 2019). The mutant was produced through combining nos-Cas9 and Obp44a-gRNA by multiple rounds of crossing and selections of the progenies using genomic PCR. The final knockout line with a 5-bp deletion in the coding region was subsequently verified via sequencing. In homozygous mutants, western blot analysis confirmed the absence of endogenous Obp44a expression.

### 3D protein structure homolog search and structure modeling

To gain insights into the Obp44a protein structure and explore potential homologous proteins, we first use I-TASSER server to search for the structure homologs of Obp44a protein: https://zhanggroup.org/I-TASSER/, which identified AeObp22. The protein alignment was performed using Clustal Omega (https://www.ebi.ac.uk/jdispatcher/msa/clustalo) and the 3D structure comparison figure (Fig. 2A) were prepared with ESPript3.0 (https://espript.ibcp.fr/ESPript/cgi-bin/ESPript.cgi) (Robert & Gouet, 2014). The 3D protein structure for AeObp22 and DmObp44a are from AlphaFold2 database (https://alphafold.ebi.ac.uk/) (Jumper et al., 2021; Mirdita et al., 2022).

### Behavioral analysis

All behavioral assessments were performed on flies selected within a 24-hour window post-eclosion and reared on standard fly medium until they reached an age range between 3 to 5 days.

Climbing Assays: 20 flies from each experimental group were introduced into a climbing vial marked with two target lines positioned at heights of 9.5 cm and 13.5 cm above the vial base. The vials were gently tapped at the initiation of the assay to ensure that all flies were situated at the bottom. Climbing assessments were carried out to determine climbing index and the percentage of flies surpassing the designated target heights (Fig. 3G and S6). Flies were allowed to move freely for a duration of 10 seconds, during which images of the climbing vials were captured to calculate the proportion of flies reaching each target height. Each experimental group underwent this procedure six times, with the final percentage representing the average of these six trials. Climbing index was computed as follows: (% of flies between 9.5 and 13.5 cm lines × 0.5) + (% of flies above the 13.5 cm line × 1). Climbing time curves were generated by allowing flies to move freely for 1 minute, capturing images of the climbing vials every 10 seconds, and calculating the proportion of flies above the 13.5 cm line at each time point.

Drosophila Activity Monitor (DAM) Recording for Sleep and Locomotion analysis: We followed the general set-up protocol, that are published (Chiu et al., 2010; Pfeiffenberger et al., 2010) and provided by the manufacturer (Trikinetics Inc.). Briefly, individual male flies were introduced into each monitor tube, containing fly food on one side and a cotton wick on the other, and placed into DAM2 monitors. The monitors were subsequently positioned within a 25 °C incubator with humidity control set to 60% relative humidity (RH) and left undisturbed for the duration of the recording. The recording schedule consisted of 3 light-dark (LD) days followed by 6 constant darkness (DD) days. Fly activity was recorded at one-minute intervals using DAMSystem311X data acquisition software. Data preprocessing was conducted utilizing the DAMFileScan113X program, followed by analysis of locomotion and sleep parameters using two MATLAB programs: the Sleep and Circadian Analysis MATLAB Program (S.C.A.M.P) (Donelson et al., 2012) and SleepMat software program (Sisobhan et al., 2022). Only data from flies that remained alive throughout the entire 9-day recording period were included in the analysis.

H_2_O_2_ Survival Probability: Following the H_2_O_2_ treatment, monitor tubes and flies were prepared in a manner akin to the procedure for DAM recordings described above, with slight modifications. Male flies were loaded into monitor tubes containing fly food with 5% H_2_O_2_ on one side and a cotton wick soaked in a 5% H_2_O_2_ solution on the other side. Recording commenced immediately after all monitors were set up, capturing fly activity at one-minute intervals over 4 LD days using DAMSystem311X software. The 1st recorded minute for analysis was the 1st minute of the subsequent hour after monitor setup (e.g., if monitor setup occurred at 5:40 PM, the initial minute for analysis would be 6:01 PM). Raw data were pre-processed using the DAMFileScan113X program and locomotion activity was analyzed using SCAMP. Survival probability was determined by checking activity levels every 6 hours, with flies considered deceased if no activity was detected within that time frame. Groups were defined as consisting of 16 flies (equivalent to half of a monitor), and multiple groups were examined for each genotype. Survival probability was calculated as the mean percentage of living flies within each group at each time point.

### tert-Butyl Hydroperoxide (tBH) Treatment

All experimental cohorts were reared on conventional fly media and collected within a 24-hour window post-eclosion. One day old flies were transferred to vials containing a medium composed of 1% agarose supplemented with 0.1% tBH and 10% sucrose, as the experimental group, while the control group were introduced to vials containing 1% agarose and 10% sucrose, both for a duration of 24 hours. Following this treatment period, adult fly brains were dissected and processed through standard immunohistochemistry procedures. For the groups with a recovery period, flies were transferred back to standard fly food after the 24-hour tBH treatment and cultured for four days before samples were collected for brain dissections.

### Lipid droplet staining

Larval or adult brains were dissected and fixed in 4% of PFA/PBS at room temperature for 30 min, and washed in PBST (PBS, 3% TritonX) three times for 20 min each time. Fixed brains were incubated in the Nile Red solution (Sigma Aldrich, 19123, diluted 1:100 in PBST from a 100 mg/ml stock solution in acetone) at 4°C overnight and then washed in PBST for 30 min each time. Larval brains were mounted on glass slides with the anti-fade mounting solution and imaged with a Zeiss LSM700 upright confocal microscope.

### Immunohistochemistry

For whole mount brain immunohistochemistry, larval and adult brains were dissected and fixed in 4% PFA/PBS at room temperature for 30-40 min, followed by washing in PBST (0.3% Triton-X 100 in PBS), and incubating in the primary antibody overnight at 4°C. On the next day, brains were washed with PBST and incubated in the secondary antibody at room temperature for 1-3 hour before final washes in PBST and mounting on the slide with the anti-fade mounting solution.

For Immunohistochemistry on acutely dissociated brain cells, third instar larvae brains were dissected and transferred to clean dish containing cold DPBS and were cut into smaller pieces by needles. After proteinase treatment (Collagenase/ Dispase [1 mg/ml] and liberase I [0.1 Wünsch units/ml] for 40 min at 25°C), media neutralization, and centrifuge, cells were resuspended in 50μl Schneider’s Insect Medium and transferred onto chambered cell culture slides (VWR, 53106-306). The chamber slides were pretreated with 0.25mg/ml Concanavalin A (Sigma). After 10min at room temperature, extra solution was removed from the slides. Adhered cells were then fixed in 4% paraformaldehyde in PBS for 10 min, then washed with PBST three times, and incubating in the primary antibody overnight at 4°C. On the next day, slides were washed with PBST and incubated in the secondary antibody at room temperature for 1 hour before final washes in PBST and mount with the anti-fade mounting solution after removal of the chamber top.

Primary antibodies used were rabbit anti-Obp44a (self-made, 1:500), mouse anti-repo (DSHB 8D12, 1:10), and mouse anti-SREBP1 (BD Pharmingen, 557036, 1:50), rat anti-elve (DSHB 7E8A10, 1:20). Secondary antibodies (1:500 dilution) used were donkey anti-rabbit Rhodamine (Jackson Immuno Research Labs, 711-295-152), goat anti-mouse Alex 647 (Invitrogen, A-32728), donkey anti-mouse CY3 (Jackson Immuno Research Labs, 715165150). Images are taken either with Zeiss LSM700 and LSM780 confocal microscopes in the lab or Zeiss LSM800 confocal microscope at NINDS Neurosciences Light Imaging Facility.

### Calcium imaging

Late 3rd instar larvae expressing *Pdf-Gal4* driving *UAS-GCaMP7f* were used for calcium imaging experiments as described (Yuan et al., 2011). Imaging was performed on a Zeiss LSM 780 confocal microscope equipped with a Coherent Vision II multiphoton laser. Larval brain explants were dissected in the external saline solution (120 mM NaCl, 4 mM MgCl_2_, 3 mM KCl, 10 mM NaHCO_3_, 10 mM Glucose, 10 mM Sucrose, 5 mM TES, 10 mM HEPES, 2 mM Ca^2+^, PH 7.2) and maintained in a chamber between the slide and cover-glass and imaged with a 40x water objective using 920 nm excitation for GCaMP signals. GCaMP7f signals were collected at 100 ms/frame for 2000 frame during each recording session. Light stimulations of 100 ms duration were delivered using a 561nm confocal laser controlled by the photo bleaching program in the Zen software. The laser power was set at 10%. GCaMP7f signals at the axonal terminal region of LNvs were recorded and analyzed. Average GCaMP7f signals of 20 frames before light stimulation was taken as F0, and ΔF (F-F0)/F was calculated for each time point. The average value of ΔF/F for individual brain samples were used to generate the average traces of calcium transients. The shaded area represents the standard-error of mean. The sample number n represents number of individual animals subjected to the optical recordings.

### Protein expression and purification

The expression and purification of ^15^N-labeled OBP44a in the apo state were carried out following the recently published protocols (He et al., 2023). The concentration was determined by the absorbance at 280 nm measured using NanoDrop Microvolume Spectrophotometer (ThermoFisher Scientific) and the predicted extinction coefficient of the protein. Of crucial importance are the use of the Shuffle *E. coli* expression host to ensure proper disulfide pairings and fold of the protein, as well as the final HPLC purification step to remove any fatty acids bound to the protein during the expression.

### NMR spectroscopy

The ^15^N-labeled OBP44a NMR samples (250 μL) were prepared by dissolving the protein in a lyophilized form in a 20 mM Potassium Phosphate buffer pH 6.6, 0.5 mM EDTA, and 10% D_2_O. The pH of each sample was checked and adjusted by the addition of 0.1N NaOH. The protein concentration used were between 70 to 400 μM in a Shigemi tube (Shigemi Co. Ltd). All fatty acids were dissolved in either DMSO (C16:0, C18:0, C18:2) or ethanol (C22:0) to create stock solutions with concentrations between 5.5 and 25 mM depending on their solubility. The final ratio of protein to fatty acid was 1:1.2 in all NMR samples.

The one-dimensional ^15^N-edited proton spectra (Bodenhausen, 1980) were acquired at 298 K on a Bruker 600 MHz spectrometer equipped with a 5 mm triple resonance probe with tri-axial gradient using a minimum of 256 scans, 2K time domain points, 8K Hz spectral width and 1 s recycle delay. The number of scans for each sample was adjusted relative to their concentrations to result in spectra of comparable signal to noise. Spectra were processed and plotted using Bruker TOPSPIN program (Bruker NMR, Billerica, MA).

### Gel-shifting assay

Fatty acid-C16 (ThemoFisher D3821) was dissolve in DMSO to make 2.1 mM stock solution. 0.5 µL of 2.1 mM FA-C16 was added to 100 µL PBS to make a final concentration of 10.5 µM. Purified OBP44a protein was added to make final concentrations of 0, 1.77 and 3.54 µM respectively. After incubating at room temperature for 40 mins in the dark, 2X Tris-Glycine native sample buffer (ThemoFisher LC2673) was added to each sample. Samples were then loaded on to two 10-20% Tris-Glycine gels (ThemoFisher XP10205BOX) in one mini gel tank and subjected to gel electrophoresis with native running buffer (ThemoFisher LC2672) in the dark. After electrophoresis, one gel was stained with 0.01% Coomassie blue R250 then scanned at 700 nm channel using Odyssey infrared scanner (Li-Cor). Another gel was directly imaged at Cy2 channel using Typhoon laser-scanner (GE Bioscience).

### Mass spectrometry and metabolomics analysis

Liquid chromatography–mass spectrometry (LC-MS/MS):

Metabolite extraction from Drosophila larval brains was executed following a modified cold methanol method as previously described (Pruvost et al., 2023). For each biological replicate, a pool of 100 brains from third instar larvae of either wild type (Canton-S) or Obp44a^-/-^ was dissected and combined in PBS buffer. Three independent biological replicates and three technique replicates were prepared for each genotype. To extract metabolites, the brain samples were homogenized in 200 μl of cold Methanol/Water solution (80/20, v/v) and subjected to gentle sonication using a Bioruptor instrument (30 s on, 30 s off, 10 cycles) at 4°C. Subsequently, the lysates were centrifuged at 10,000 x g for 10 minutes at 4°C, and the resulting supernatants were collected for LC-MS/MS analysis. LC-MS/MS experiments were performed utilizing a combination of ZIC-HILIC chromatography with an Acetonitrile/Water/7mM Ammonium acetate solvent system, coupled with high-resolution mass spectrometry. The analysis was conducted using a maXis-II-ETD UHR-ESI-Qq-TOF mass spectrometer (Bruker Daltonics) equipped with a Dionex Ultimate-3000 liquid chromatography system. The ZIC-HILIC column (2.1mm x 150mm) operated under mildly acidic pH conditions enabled efficient separation of the target metabolites. Global metabolic profiling was achieved by conducting three technical replicates for each of the three biological replicates per genotype, ensuring robust and comprehensive metabolite analysis.

LC-MS/MS metabolomics analysis:

Metabolomic analysis of samples was performed using MetaboScape-2023 and MetaboAnalyst software, following established protocols (Xia & Wishart, 2011). Compound identification based on the Bruker MetaboBase Personal 3.0, MoNA, MSDIAL, METLIN, and HMDB97 metabolomic libraries resulted in the annotation of 1371 out of the total 3140 features detected. To evaluate data quality and variation, Principal Component Analysis (PCA) was conducted on the 1371 annotated features. A 3D plot was generated using Omicshare tools (https://www.omicshare.com/tools). Accurate mass measurements, with an accuracy of less than 5 parts per million (ppm), and MS/MS spectra were both utilized for robust metabolite and lipid identification. In total, 300 metabolites were identified to be present in both wild-type and mutant samples. Statistical significance between the three biological replicates of wild-type and mutant animals was assessed using Student’s t-test. Metabolites with a p-value less than 0.05 were considered significant. The data were visualized using various tools. A volcano plot was generated to illustrate the differential expression of metabolites using TBtools (Chen et al., 2020). Heatmaps of changed metabolites were also constructed using TBtools. The metabolomic datasets generated in this study will be deposited in the National Metabolomics Data Repository (NMDR) and will be made publicly accessible upon the publication of the study.

### Western Blot

To test Obp44a level in mutant and wild type, five 3^rd^ instar larval brains were dissected and homogenized in 30 μl lysis buffer (N-Per lysis buffer (Thermo Fisher, 87792) containing 0.1 mM DTT and 1:100 diluted proteinase inhibitor cocktail (Sigma, P8340)). To test the secretion of Obp44a, five Obp44a-GFP-KI 3^rd^ larval brains were dissected and introduced into 25ul external saline solution (120 mM NaCl, 4 mM MgCl2, 3 mM KCl, 10 mM NaHCO3, 10 mM Glucose, 10 mM Sucrose, 5 mM TES, 10 mM HEPES, 2 mM Ca2+, PH 7.2) for 0 min, 30 min and 60 min. The solution was recovered, and the remaining brains were homogenized in 30 μl lysis buffer. Larval hemolymph samples were collected from third instar larvae using a method described in an online video resource (https://www.youtube.com/watch?v=im78OIBKlPA), with twenty larvae used to obtain hemolymph for each genotype. The supernatant from brain lysis was incubated with protein sample buffer containing 10 mM DTT and heated to 95°C for 5 min. Hemolymph samples were incubated with protein sample buffer containing 10 mM DTT at room temperature for 30 minutes. The prepared protein samples were loaded onto an SDS-PAGE gel. Subsequently, the separated proteins were transferred onto a PVDF membrane. To block nonspecific binding, the membrane was treated with a 5% milk solution in TBST. Following blocking, the membrane was incubated overnight at 4°C with primary antibodies, including rabbit anti-Obp44a (1:5000) or rabbit anti-GFP (Abcam, ab6556, 1:2000), or rabbit anti-alpha tubulin (Abcam, ab15246, 1:2000). After three washes with TBST, a secondary antibody (HRP-conjugated) was applied at a dilution of 1:10,000 and incubated for 1 hour at room temperature. Membrane exposure was performed using a chemiluminescence detection kit (Bio-Rad, 1705062).

### DHE staining

To assess superoxide levels in live brain samples, we employed Dihydroethidium (DHE, Life Technologies, D11347) staining and live *ex vivo* imaging, following a protocol adapted from Bailey et al. (Bailey et al., 2015). Early third instar brains were dissected in Schneider’s medium and incubated in a 30 μM DHE solution in a dark environment for 30 minutes. Subsequently, the brains underwent two washes in Schneider’s medium before being mounted in an external saline solution, facilitating immediate confocal imaging. It’s important to note that all brain samples were freshly prepared and imaged to ensure the reliability of the results.

### In vivo imaging of Obp44a release with C12-red fatty acid feeding

Obp44a::GFP knock-in larvae were transferred to food containing 1μg/ml C12-red fluorescent fatty acids (BODIPY™ 558/568 C12, Thermo Fisher, D3835) at 72h after fertilization. After feeding for 24h, larval brain was dissected in external saline solution (120 mM NaCl, 4 mM MgCl2, 3 mM KCl, 10 mM NaHCO3, 10 mM Glucose, 10 mM Sucrose, 5 mM TES, 10 mM HEPES, 2 mM Ca2+, PH 7.2) and maintained in a chamber between the slide and cover-glass and imaged with a 40x objective using Zeiss 800 confocal microscope or Nikon A1R confocal microscope for 30-90 minutes.

### Quantification and statistical analysis

For lipid droplet density quantification, images were processed by Imaris 3D image visualization software. The Spots module of the Imaris was used to detect the number of lipid droplets within a 3D volume of 88.7 µm x 88.7 µm frame with thickness of 17 µm in larval brain (Fig. 3D). The total number was exported from Imaris into Excel. The density was calculated by lipid droplets numbers divided by the volume of quantified region. For adult brain (Fig. 3E), lipid droplets number was manually calculated within the optic lobe OLC region with the thickness covered OLC. The total numbers in OLC were used for statistical analysis.

*In vivo* glutathione redox (GSH/GSSG) biosensor (mito-roGFP2-Grx1) redox level (Fig. 4F-G) in 10-day old flies was calculated by the mean intensity of 405nm channel divided by mean intensity of 488nm channel in the whole optic lobe region with a frame of 15.8 µm x 15.8 µm (Albrecht et al., 2011). For each brain optic lobe sample, the top, median and bottom three sections were used for quantification. A ratio image was created by dividing the 405 nm image by the 488 nm image pixel by pixel and displayed in false colors using the lookup table ‘‘Fire’’ in ImageJ.

The superoxide radical level revealed by DHE staining (Fig. S7) was quantified by the mean intensity for the red channel in the individual astrocyte nucleolus region.

Graphing and statistics analysis of the quantifications were performed using Graphpad Prism (9.4.1) For statistical analyses between two groups of samples, two-tailed unpaired student *t*-test (unpaired t-test with Welch’s correction) was performed; for experiments with more than two groups, one-way ANOVA analysis followed by multiple comparisons Tukey post-hoc test or Welch’s ANOVA test with Dunnett’s multiple post-tests was performed. Exact value of sample number n, the statistical testes used and the confidence intervals and the precision measures for individual experiments are included in the figure legends. Most quantitative data are presented as bar plot overlaid with dot plot; bar plot shows the mean (height of bar) and SEM (error bars); dot plot displays individual data points. n represents groups with 20 flies per group in Fig. 2A and Fig. S4. n represents the fly numbers in Fig. 2C, 2D. n represents groups with 16 flies per group in Fig. 4E. n represents number of larvae brains in Fig. 2H, 3D, 3F, 3G. n represents number of adult brains in Fig. 3E. n represents number of quantified sections in 4F, 4G (3 sections per brain). n represents number of quantified cells in Fig. 3I, 3J, S5B. Statistical significances were assigned as: *: *p* < 0.05, **: *p* < 0.01, ***: *p* < 0.001, ns: not significant.

Schematic images in Fig. 2C, 3B, 4A, 5B, 5H, and fig. S4 are created with BioRender.com.

---

**Figure S1.**
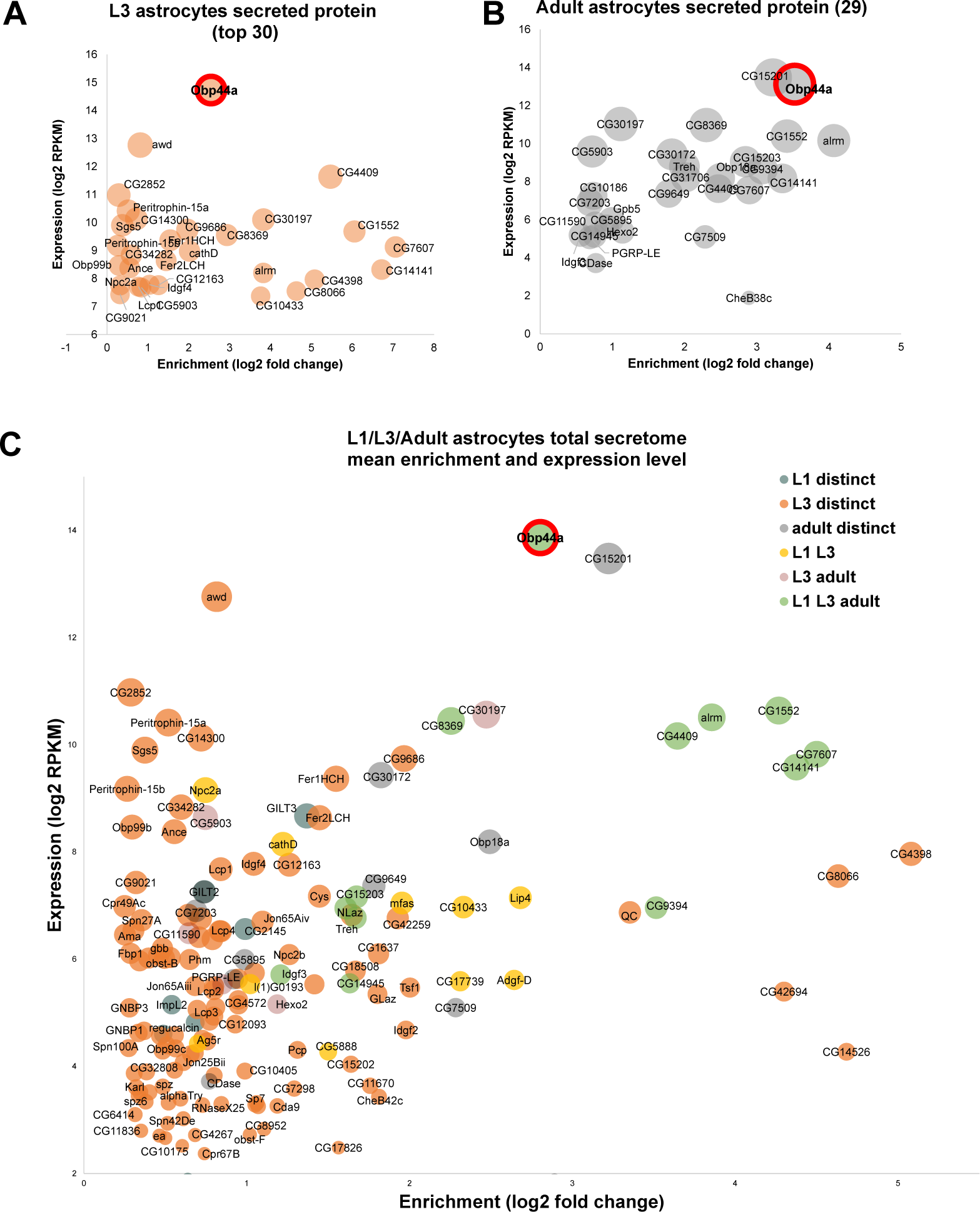
Astrocyte-secreted molecules during development. (A) The top 30 highly expressed and enriched astrocyte-secreted proteins identified in 3rd instar (L3) larval astrocytes. (B) The 29 highly enriched astrocyte-secreted proteins identified in adult astrocytes. The data are generated using published RNAseq datasets by Huang Y et al. (C) Expression level and enrichment analysis of 156 enriched astrocyte-secreted molecules across three developmental stages, including L1, L3, and adult stages. 13 secreted proteins are found in all three stages (green circles), with Obp44a being the most abundantly produced secreted protein in astrocytes. Data cutoff: enrichment log2 fold change ≥ 0.25 & RPKM ≥5.

**Figure S2.**
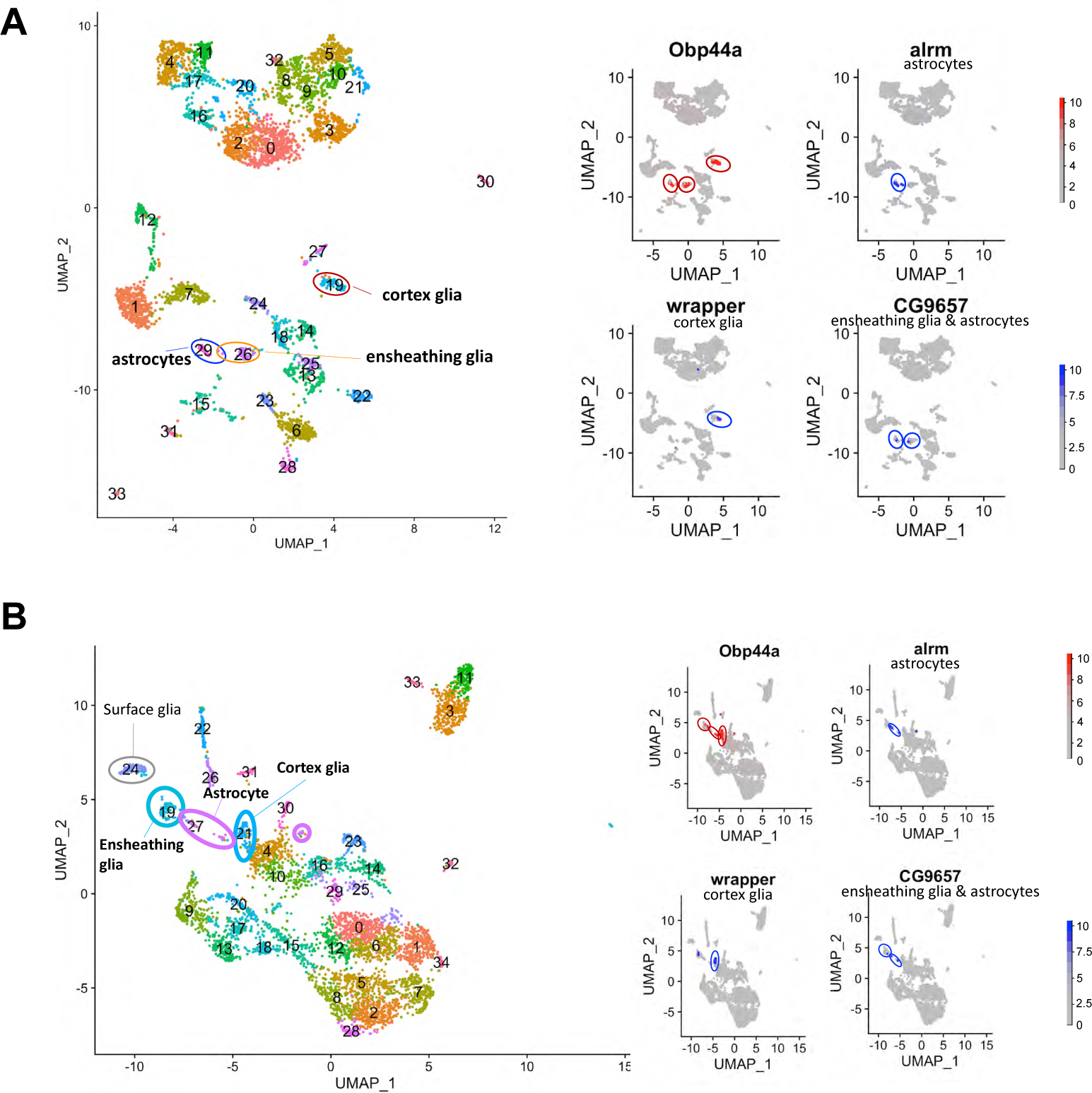
Obp44a is enriched in astrocytes, cortex and ensheathing glia in larval brains. (A) Cell atlas of the first instar (L1) larval brain. (B) Cell atlas of the 3rd instar (L3) larval brain. Obp44a exhibits high expression in three distinct cell clusters: astrocytes, ensheathing glia, and cortex glia. Corresponding markers for each glial cell type (alrm, wrapper, and CG9657) are displayed in the right panel. The data are generated using published RNAseq datasets by Avalos CB et al. and Ravenscroft TA et al.

**Figure S3.**
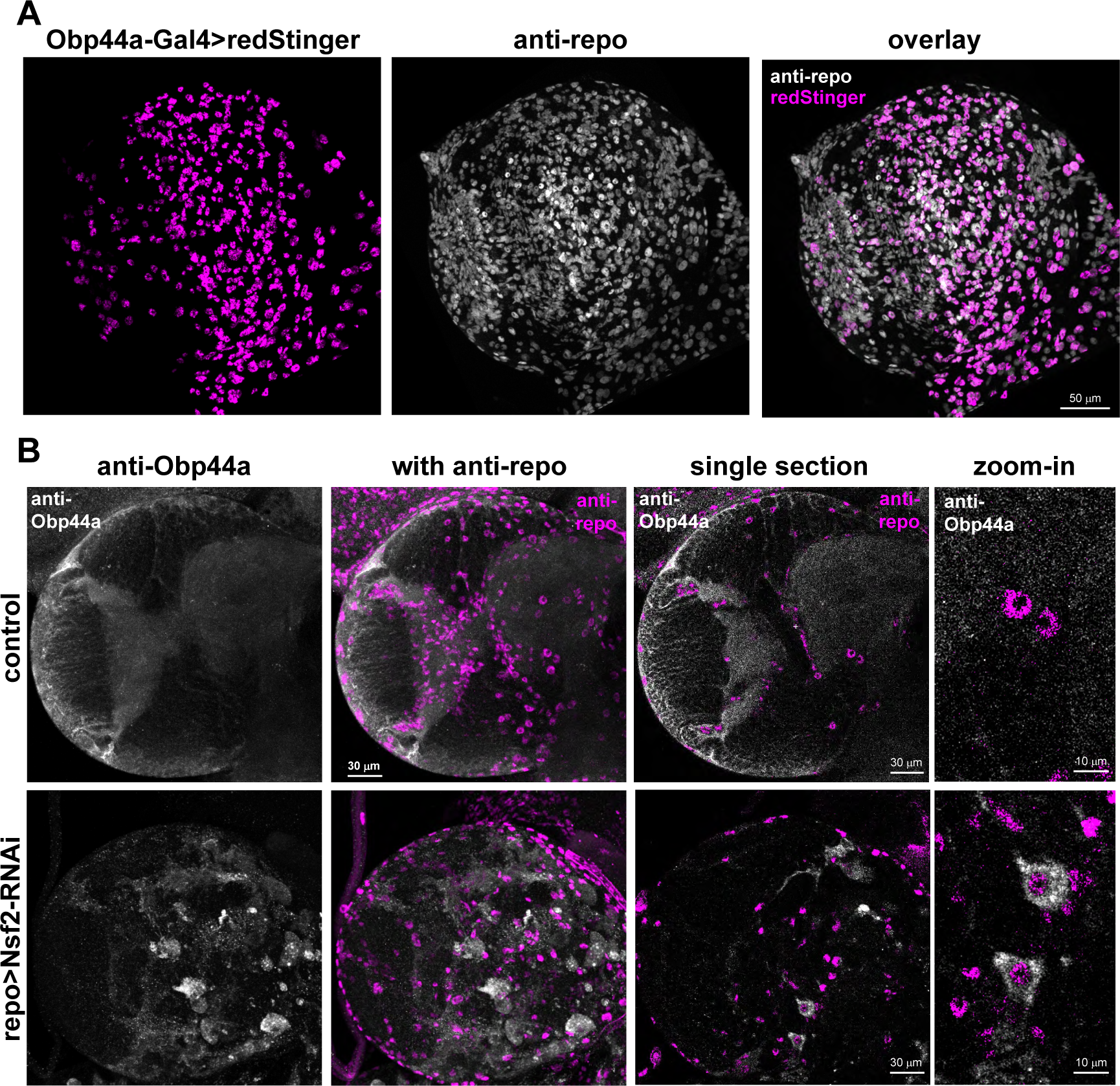
Obp44a is a glia derived secreted protein. (A) Obp44a-Gal4 driven redStinger is colocalized with repo-positive nuclei, indicating Obp44a is expressed in glia cells. (B) Inhibiting glia secretion using Repo-Gal4 driven Nsf2 RNAi effectively blocks Obp44a secretion, leading to its cumulation within the soma of repo-positive glial cells (bottom), in contrast to the diffuse distribution of the control group (top). Representative MIP confocal images of the 3rd instar larval brains are shown. The immunostaining experiments are performed using anti-Obp44a (grey) and anti-Repo (magenta) antibodies.

**Figure S4.**
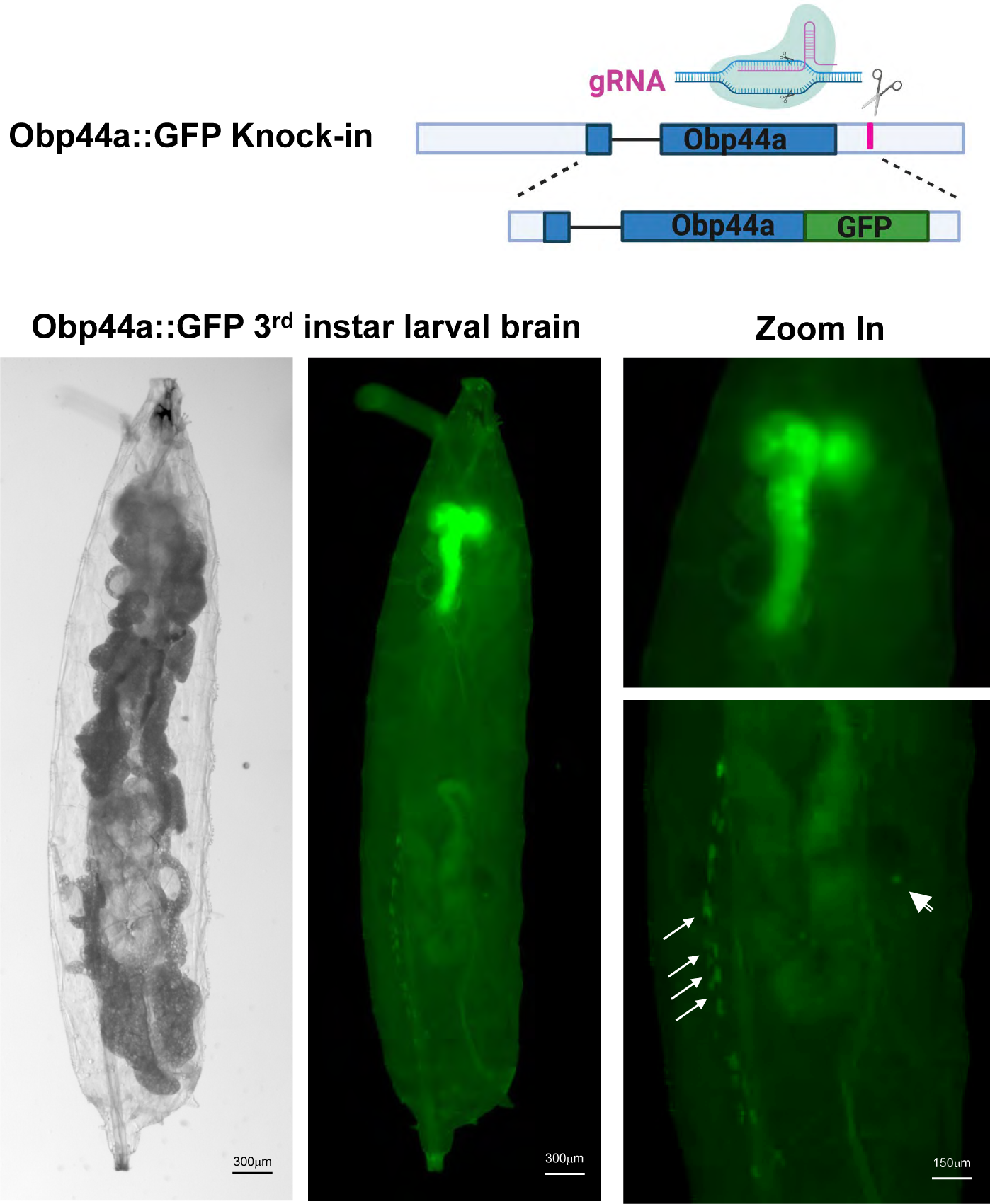
Obp44a is a glia derived secreted protein concentrated in the fly CNS. (A) Schematic diagram illustrating the CRISP-Cas9 mediated gene editing strategy used to generate the knock-in Obp44a::GFP line (top). (B) The distribution of Obp44a::GFP is concentrated in the CNS. Left: the transmission light image showing a 3^rd^ instar larva. Right: Only the larval brain (top) and nephrocytes on the larval body wall (arrow), and a part of the male testis (arrowhead) show clear GFP signals. Similar to anti-Obp44a staining, the signal in the larval brain prominently localizes in the neuropil regions.

**Figure S5.**
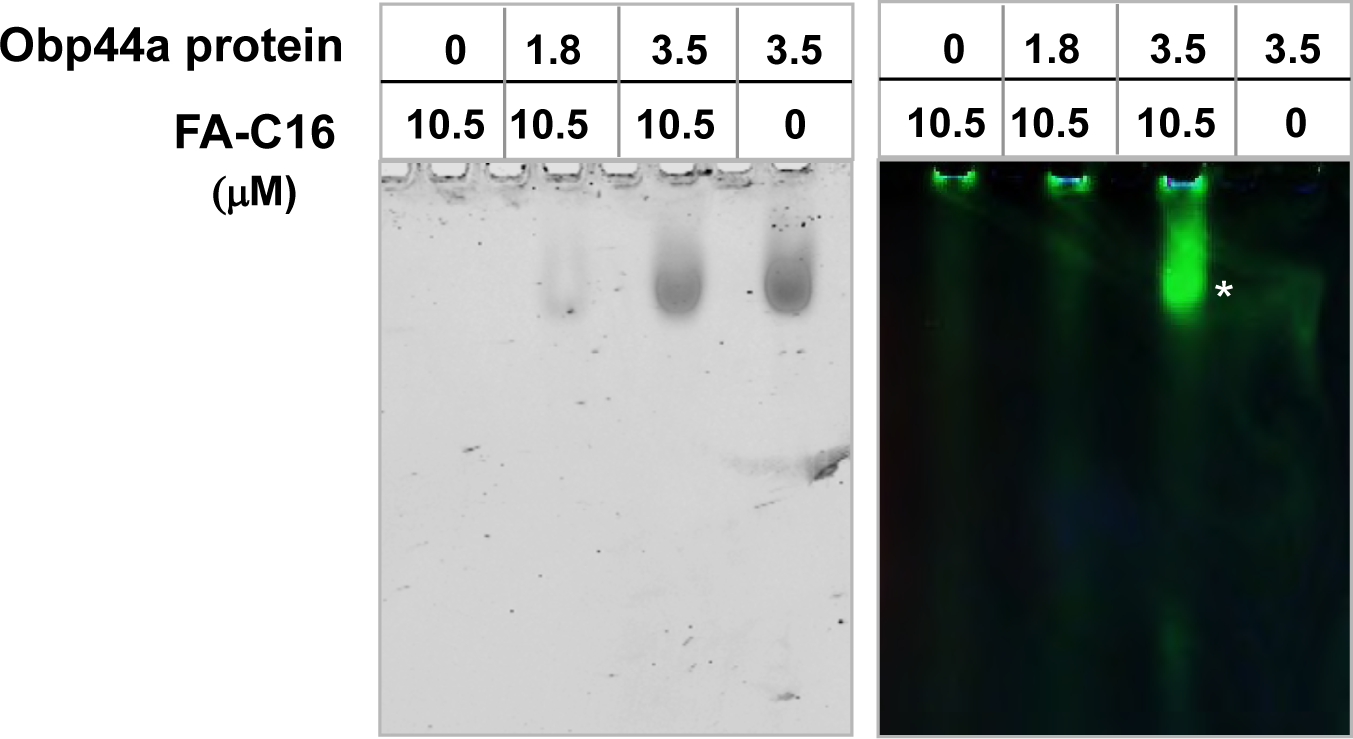
Obp44a binds fatty acids. Native PAGE binding assay demonstrating the interaction between Obp44a protein and fluorescent fatty acid C16. Coomassie-stained purified Obp44a protein is shown on the left, while the right panel displays BODIPY FL-labeled green fluorescent C16 fatty acid (FA-C16). FA-C16, when alone, remains at the well without migrating along the lane. Upon the addition of 3.5 μM of OBP44a to 10.5 μM of FA-C16, migration occurs, and colocalization with Obp44a (indicated by a white star) suggests binding of Obp44a to FA-C16.

**Figure S6.**
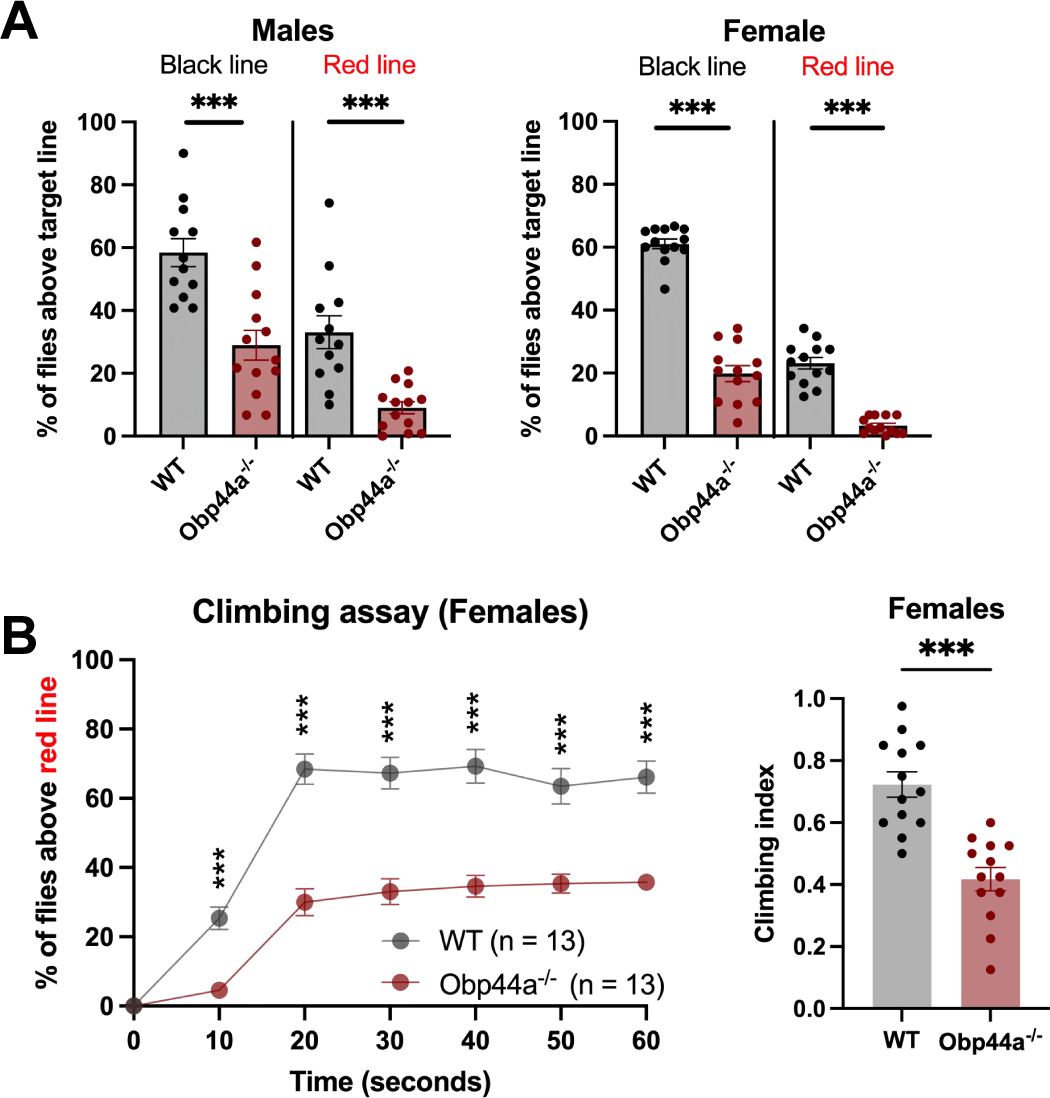
Reduced climbing ability in Obp44a^-/-^ flies. (A) Percentiles of male and female flies capable of climbing beyond the indicated black and red lines (Figure 2A). (B) Changes in climbing percentiles and climbing index of female flies over time. Statistical significance was determined using a two-tailed Student’s t-test. ***P<0.001. Error bars represent mean ± SEM; n=12,13 groups. Ach group contains 20 flies.

**Figure S7.**
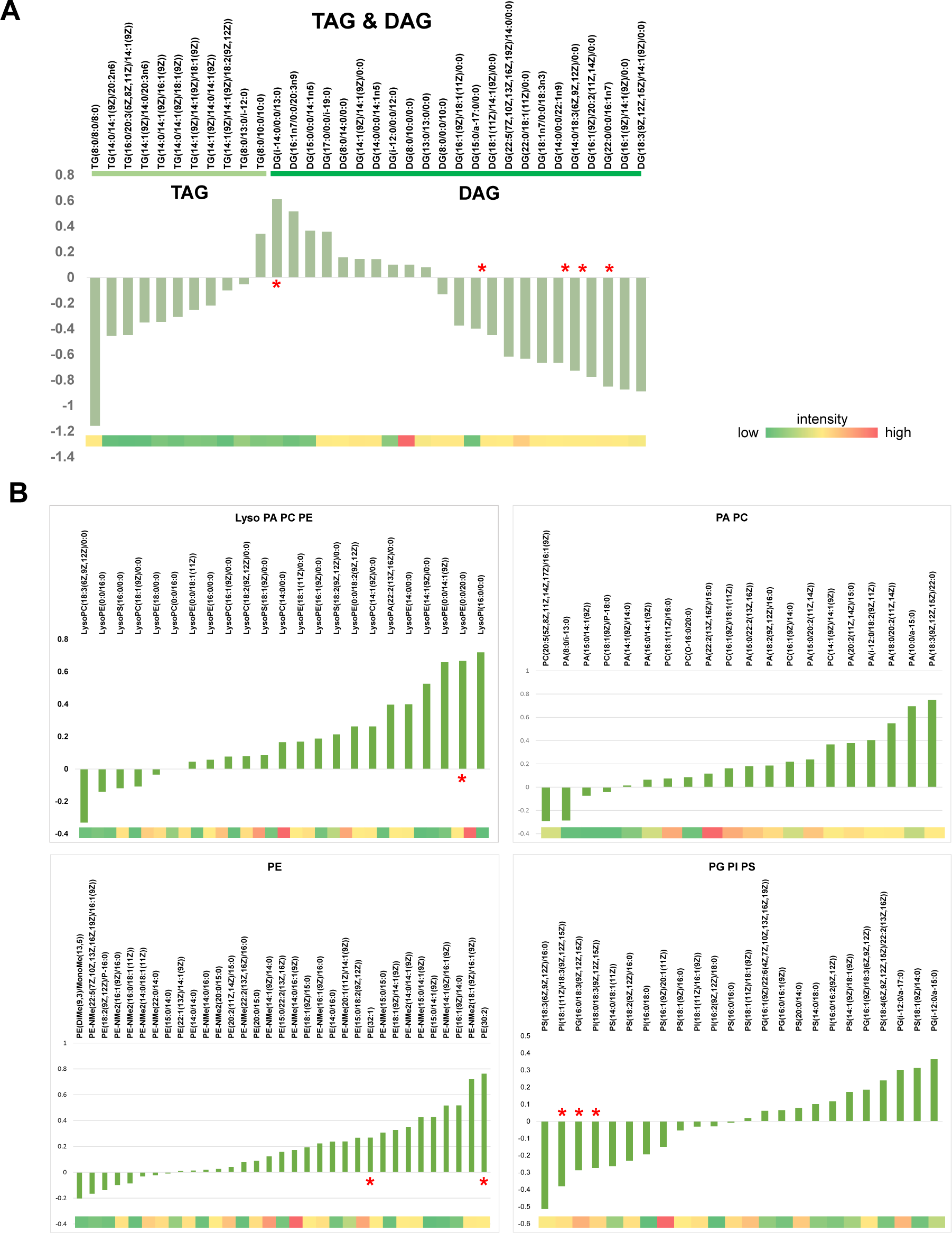
Brain metabolomic alterations in Obp44a^-/-^ mutants. (A) Alterations in the levels of all detected TAG and DAG. (B) Alterations in the levels of all detected phospholipids. Metabolites marked with a red star have p-values<0.05.

**Figure S8.**
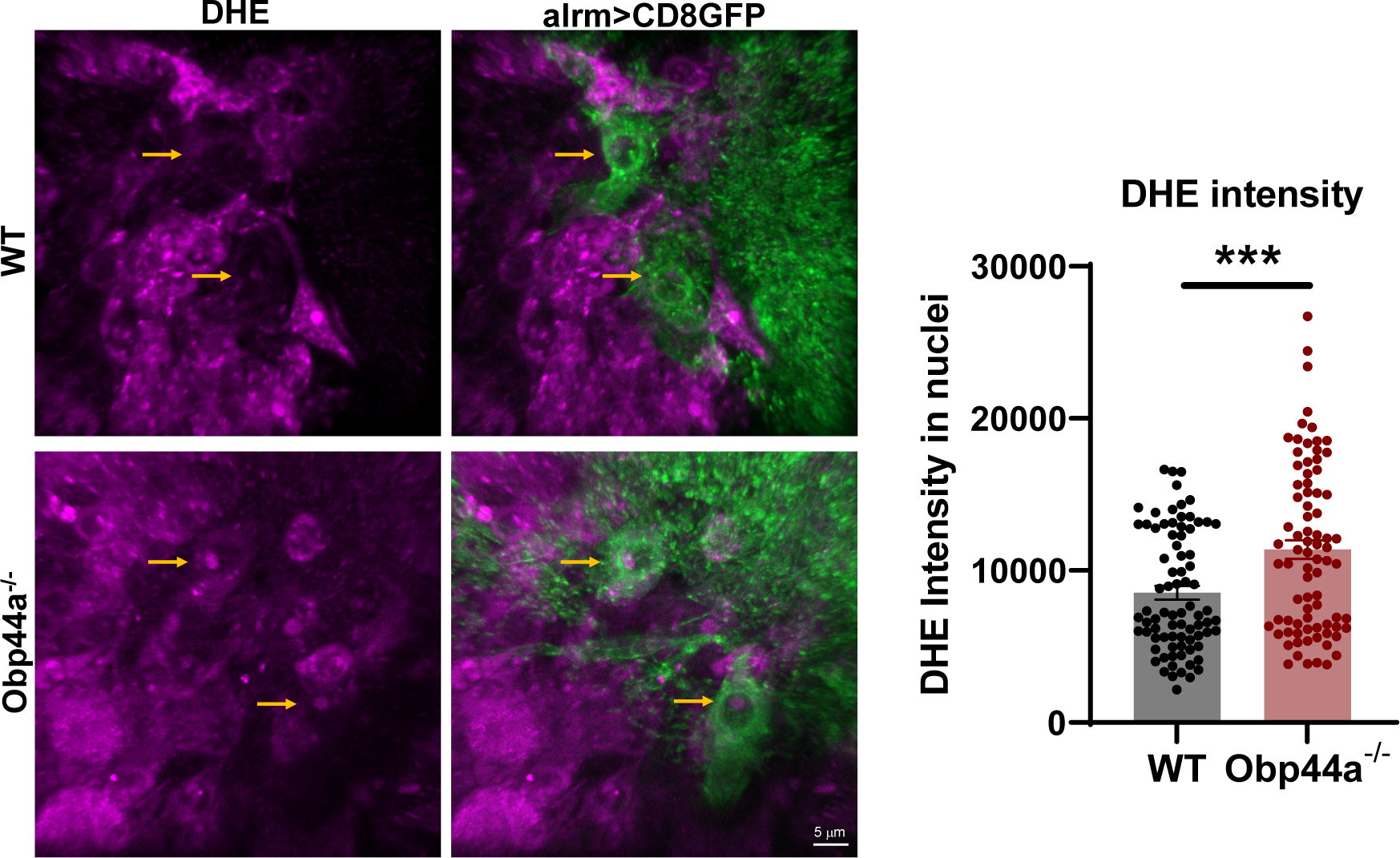
Increased superoxide radicals in Obp44a^-/-^ mutant brain by Dihydroethidium (DHE) staining, which reveals superoxide radical levels in nuclei. DHE intensity was quantified in astrocyte nuclei within the larval brain. Statistical significance was determined using a two-tailed Student’s t-test. ***P<0.001. Error bars represent mean ± SEM; n=80, 81.

**Figure S9.**
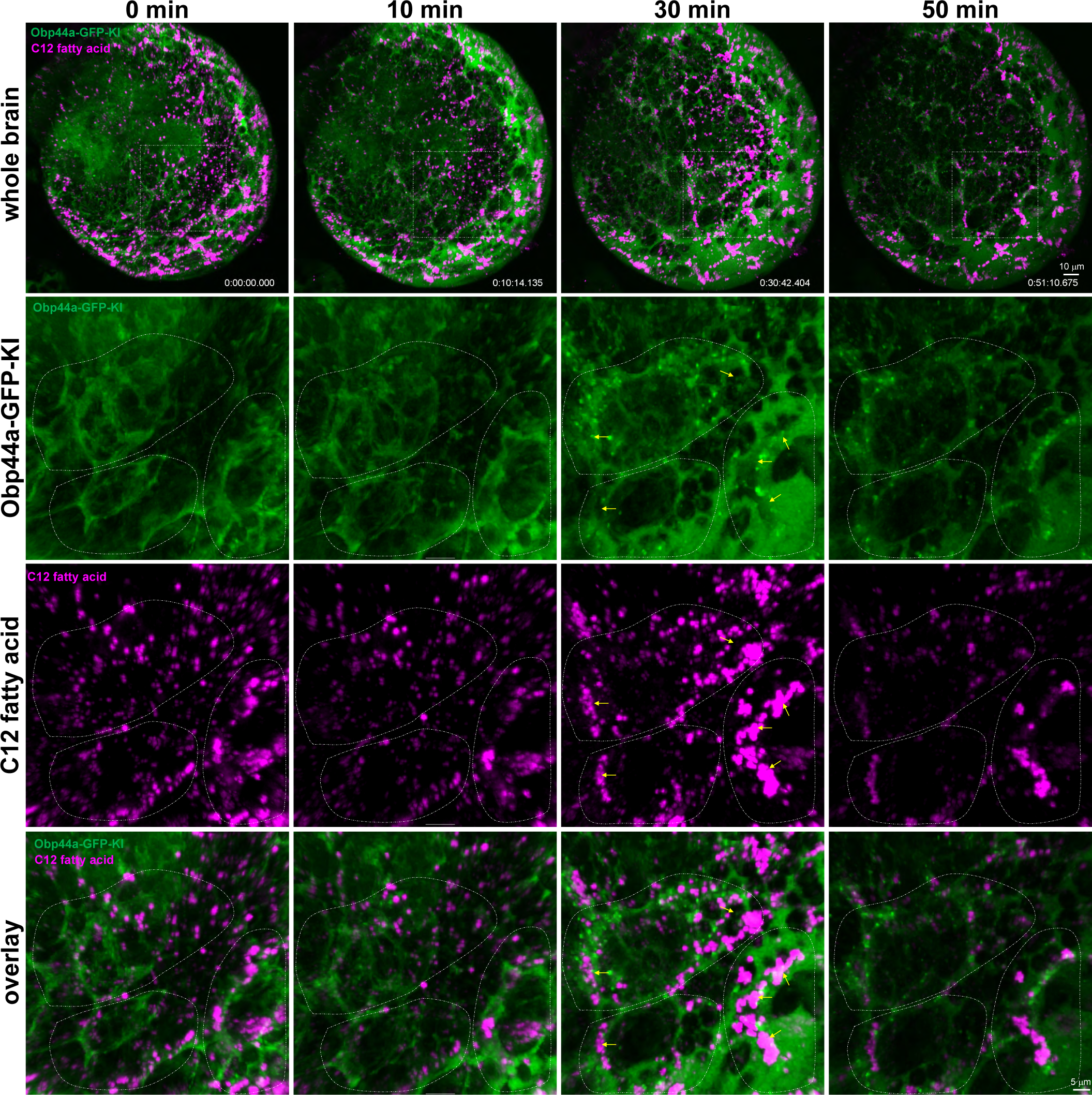
Obp44a mobilizes fatty acid cargos *in vivo*. Timelapse *in vivo* imaging with zoom-in depicts the shift of Obp44a::GFP and C12-red localization from the central to the surface brain. At 30 mins, C12-red fatty acids are mobilized to the ensheathing glia region near surface brain as big particles (yellow arrows) with Obp44a wave and diminished at 50 mins when Obp44a is released out of the brain.

**Figure S10.**
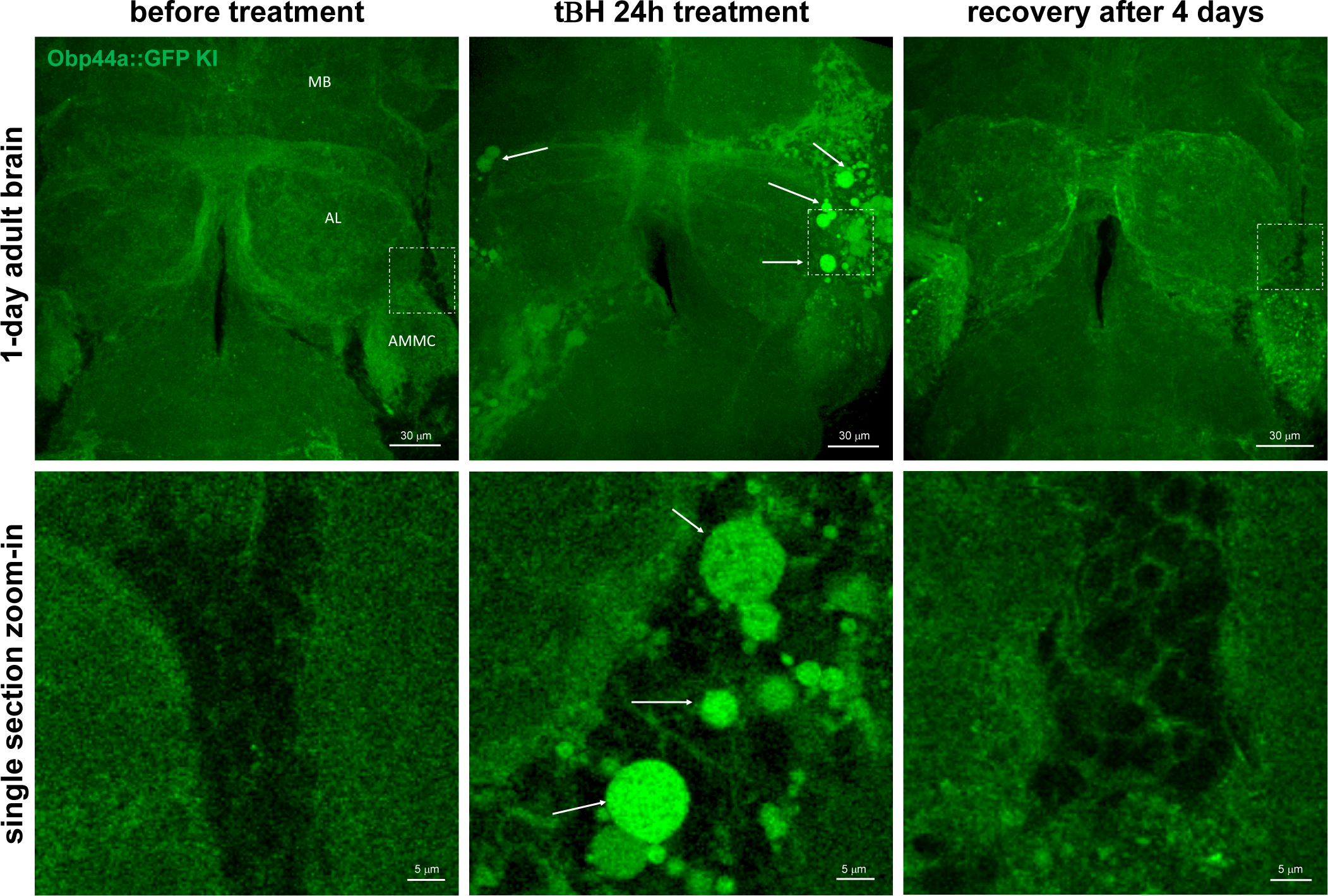
Obp44a response to severe oxidative stress. The distribution of OBP44a::GFP in the adult brain altered following a 24-hour exposure to tert-butyl hydroperoxide (tBH), a known inducer of reactive oxygen species (ROS) and oxidative stress. Obp44a-GFP exhibited rapid accumulation as large intracellular droplets (indicated by arrows) in specific brain regions, including the mushroom body (MB), antenna lobe (AL), and antennal mechanosensory and motor center (AMMC) area. This aberrant distribution pattern reverted to normal after four days of incubation with regular food. Arrows indicate the accumulation of Obp44a droplets in response to heightened oxidative stress.

**Figure S11.**
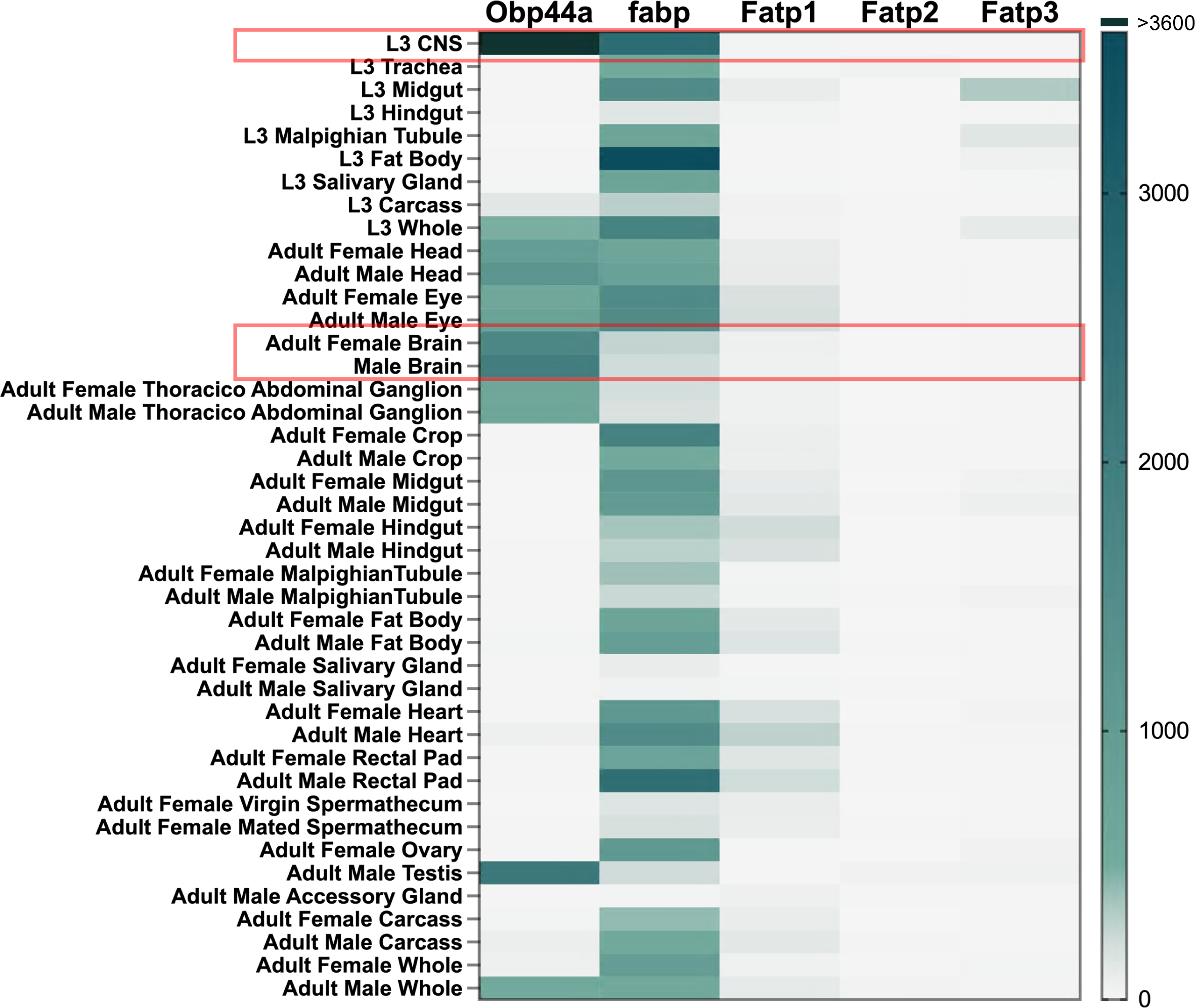
Comparison of the expression of five Drosophila fatty acid binding proteins in various tissues, highlighting Obp44a as the primary fatty acid chaperone in the brain. Obp44a exhibits high expression levels in both the central and peripheral nervous systems, including the brain, head, eye, and thoracicoabdominal ganglion. In contrast, Fabp displays a broader expression profile across various tissues, with lower expression in the brain compared to Obp44a. The three Fatp, on the other hand, exhibit notably lower expression levels across all tissues when compared to both Obp44a and Fabp.

